# Oncogenic ERK signaling represses chaperone-mediated autophagy through transcriptional control of LAMP-2A

**DOI:** 10.64898/2026.03.03.709259

**Authors:** Eva Berenger, Merve Kacal, Alice Maestri, Elena Kochetkova, Boxi Zhang, Suresh Sajwan, Mattias Mannervik, Erik Norberg, Vitaliy O. Kaminskyy, Helin Vakifahmetoglu-Norberg

**Affiliations:** Department of Physiology and Pharmacology, Solnavägen 9, Biomedicum, Karolinska Institutet, 171 65, Stockholm, Sweden; Division of Cardiovascular Medicine, Department of Medicine Solna, Karolinska Institutet, 171 64 Stockholm, Sweden; Center for Molecular Medicine, Karolinska Institutet, 171 76 Stockholm, Sweden; Department of Molecular Biosciences, The Wenner-Gren Institute, Stockholm University, 10 691 Stockholm, Sweden

**Keywords:** Cancer, Chaperone-mediated autophagy, LAMP-2A, MEK-ERK signaling, GSK-615

## Abstract

Chaperone-mediated autophagy (CMA) is a selective lysosomal degradation pathway governed by the rate-limiting receptor LAMP-2A and increasingly implicated in cancer. However, the oncogenic circuits that enforce CMA repression and whether this state is therapeutically reversible remain unclear. Here, we developed a quantitative bioluminescence-based reporter to measure CMA activity in human cancer cells and combined parallel chemical and genome-scale CRISPR-Cas9 screens to define regulatory pathways. The chemical screen identified GSK1059615 as a CMA-restoring compound that increased LAMP-2A transcription and protein abundance *in vitro* and *in vivo*. In parallel, the CRISPR screen revealed ERK signaling as a pathway-level suppressor of CMA. Genetic or pharmacologic ERK inhibition de-repressed LAMP-2A expression, while integrated modulation of ERK, PI3K-AKT, and p38 signaling coordinated transcriptional induction and stabilization of LAMP-2A. Transcriptomic analyses further implicated FOXO1/FOXP1-driven programs in LAMP-2A regulation. Together, these findings position CMA as an integrated output of oncogenic signaling networks and establish a mechanistic framework for restoring CMA activity in defined cancer contexts.

## Introduction

Chaperone-mediated autophagy (CMA) is a selective lysosomal degradation pathway specialized in the removal of proteins in a motif-dependent manner (Kaushik & Cuervo, 2018; Kirchner *et al*, 2019; Seike *et al*, 2024). CMA is typically engaged to maintain proteostasis under conditions of cellular stress (Yao & Shen, 2023), during which substrate proteins are recognized by HSC70 and delivered to lysosomes for translocation via the lysosomal-associated membrane protein 2A (LAMP-2A) (Ikami *et al*, 2022; Qiao *et al*, 2023). Among known regulatory components, LAMP-2A constitutes the rate-limiting determinant of CMA activity, with its expression, stability, and multimerization at the lysosomal membrane directly controlling CMA flux (Hubert *et al*, 2022; Qiao *et al*., 2023; Yao & Shen, 2023).

Accumulating evidence establishes CMA as a critical regulator of cancer cell proteostasis, metabolism and immune evasion (Arias & Cuervo, 2020; Hao *et al*, 2019; Kacal *et al*, 2021; Li *et al*, 2025). Notably, its role in tumor biology is highly context-dependent, functioning either as a tumor suppressor or facilitator of malignant progression depending on cancer type, stage and microenvironmental cues (Arias & Cuervo, 2020; Hubert *et al*., 2022; Liu *et al*, 2023; Rios *et al*, 2020; Saberiyan *et al*, 2025). Whereas early studies emphasized tumor-promoting roles of CMA linking oxidative stress tolerance and metabolic fitness (Chen *et al*, 2023; Ding *et al*, 2016; Li *et al*., 2025; Saha, 2012; Yan *et al*, 2024), subsequent work revealed declined CMA activity in multiple cancers, supporting a tumor-suppressive function through restriction of oncogenic protein accumulation (Gomes *et al*, 2017; Hao *et al*., 2019; Vakifahmetoglu-Norberg *et al*, 2013; Xia *et al*, 2015; Zhou *et al*, 2025). We recently demonstrated that CMA directly modulates the metastatic phenotype of advanced tumors (Zhou *et al*., 2025), showing that loss of human LAMP-2A reprograms mesenchymal cancer cells toward a dedifferentiated and aggressive state, thereby conferring metastatic competence. Consistently, CMA downregulation was observed in metastatic cancers, accompanied by suppressed LAMP-2A expression in patient tumors. Together, these observations suggest that CMA may be selectively repressed at specific disease stages and raise the possibility to achieve therapeutic benefits by restoring CMA activity. However, despite this growing rationale, pharmacological modulation of CMA in tumors remains limited, and no clinically viable strategies currently exist (Arias & Cuervo, 2020; Hubert *et al*., 2022). Whether oncogenic signaling actively imposes a reversible brake on CMA in defined tumor contexts remains unclear.

One major barrier to therapeutic targeting of CMA is the incomplete understanding of how LAMP-2A itself is regulated in cancer. To date, there is limited evidence defining signaling pathways that repress LAMP-2A expression in human cancer cells, thereby dampening CMA. Prior work identified a lysosomal mTORC2/AKT/GFAP axis that restraints CMA by destabilizing LAMP-2A translocation complex (Arias *et al*, 2015), and pharmacological inhibition of class I PI3K or FLT3 has been reported to enhance CMA activity (Endicott *et al*, 2020; Xia *et al*., 2015). However, these studies do not address cancer-context-specific transcriptional control of *LAMP-2A*. At the transcriptional level, RARα signaling, NRF2 and NFAT activation have been implicated in regulating *LAMP-2A* (Anguiano *et al*, 2013; Pajares *et al*, 2018; Valdor *et al*, 2014), but these findings largely derive from aging, stress, or immune-related settings rather than oncogenic signaling contexts (Bourdenx *et al*, 2021; Gomez-Sintes *et al*, 2022; Huang *et al*, 2025; Ueda *et al*, 2022; Valdor *et al*., 2014). Likewise, post-translational regulation of LAMP-2A, such as p38 MAPK-mediated phosphorylation, has been described primarily in neuronal systems or under ER stress (Li *et al*, 2017; Motomura *et al*, 2024). Thus, how oncogenic signaling networks converge on LAMP-2A to regulate CMA in human cancers remains largely unexplored and represents a critical gap addressed in this study.

Here, we report a quantitative bioluminescence-based reporter assay to monitor CMA activity human cancer cells. Using this platform, we conducted a small-molecule screen and identified GSK1059615 (GSK-615) as a pharmacological activator of CMA that induces transcriptional upregulation and stabilization of LAMP-2A, enhancing CMA-dependent substrates degradation without engaging macroautophagy. In parallel, we uncovered the MAPK/ERK pathway by a genome-wide CRISPR-Cas9 screen, as a previously unrecognized negative regulator of CMA in cancer cells. Mechanistically, GSK-615 restored CMA activity through coordinated inhibition of ERK signaling, relieving transcriptional repression of *LAMP-2A*, together with AKT inhibition and p38 activation, which supported LAMP-2A protein stability. Specifically, (i) ERK inhibition promoted *LAMP-2A* transcription through involvement of Forkhead family transcription factors, (ii) AKT inhibition relieved the lysosomal AKT/GFAP brake on LAMP-2A, and (iii) p38 phosphorylation further supported LAMP-2A protein stability. This integrated signaling response elevated CMA activity both *in vitro* and in xenograft tumors. Collectively, these findings define oncogenic ERK signaling as a pathway-level brake on CMA, elucidate how multiple signaling inputs converge on LAMP-2A, and establish a mechanistic framework for restoring CMA activity in cancer cells.

## Results

### Development of a bioluminescent reporter to quantify CMA in cells

To enable rapid and quantitative detection of changes in CMA activity, we engineered NanoLuc (NLuc)-based reporters that provide an inverse readout of CMA flux in living cells. Because CMA selectively degrades proteins bearing KFERQ-like targeting motifs, we fused the validated CMA substrate hexokinase-2 (HK2) (Xia *et al*., 2015) to NLuc, with or without a GFP tag, under a constitutive CMV promoter (Fig. EV1A). In this configuration, luminescence scales with reporter abundance, such that CMA-mediated lysosomal degradation of HK2 results in proportional decrease in NLuc signal, enabling a direct and plate-compatible measurement of CMA activity (Fig. 1A). To contextualize our model system, we first profiled baseline LAMP-2A protein levels across normal lung fibroblasts (WI-38, MRC-5), epithelial cancer cells (A549), and mesenchymal cancer cells (SUM159, ES2). LAMP-2A expression was lesser in cancer cells relative to normal fibroblasts and was lowest in mesenchymal cancer cell lines (SUM159, ES2) (Fig. EV1B). Notably, WI-38 and MRC-5 fibroblasts despite being of mesenchymal lineage, maintain high LAMP-2A level, indicating that LAMP-2A repression is associated with malignant mesenchymal cancer state rather than mesenchymal lineage itself, highlighting a tumor-specific constrain on CMA. Based on this rationale, SUM159 (triple-negative breast cancer) and ES2 (ovarian carcinoma) cells, representing distinct tissue origins while sharing a mesenchymal state, were used for further reporter validation and mechanistic studies (Fig. EV1C).

**Figure 1.**
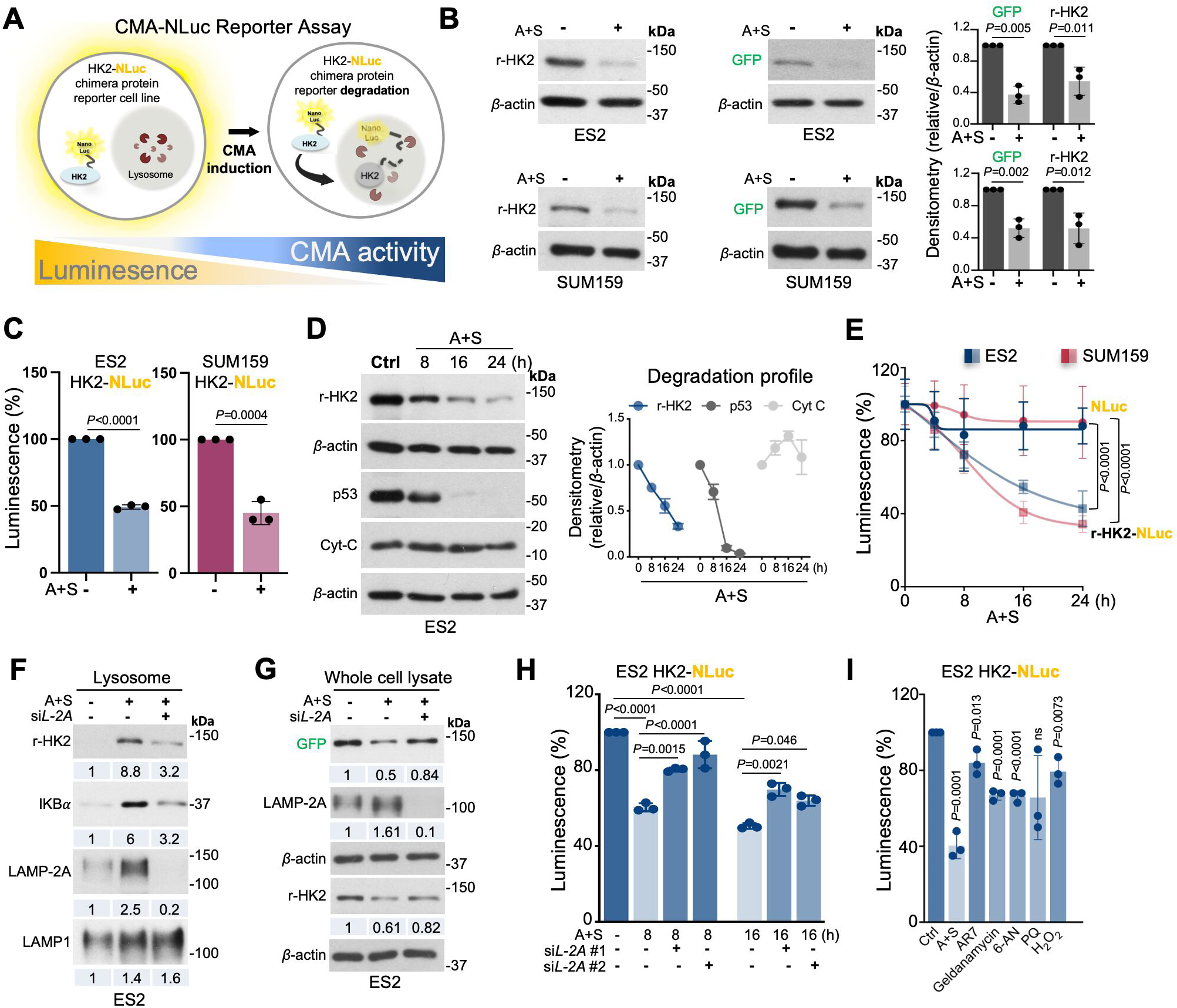
Development of a quantitative luminescence reporter to measure CMA activity. (**A**) Schematic illustration of the NanoLuc (NLuc)-based CMA reporter assay. (**B**) Immunoblot analysis of HK2-NLuc reporter degradation upon CMA activation in ES2 and SUM159 cells stably expressing the reporter treated with AC220 + Spautin-1 (A+S) for 16 h. Left and middle panels show representative immunoblots probed with anti-HK2 and anti-GFP antibodies; right panel shows quantification normalized to *β*-actin. *P* values were calculated using unpaired two-tailed Student’s *t*-test. (**C**) Relative NLuc luminescence in ES2 and SUM159 reporter cells following CMA activation with A+S for 16 h. *P* values were calculated using unpaired two-tailed Student’s *t*-test. (**D**) Time-course immunoblot analysis of reporter HK2 (r-HK2), endogenous p53 (CMA substrate) and cytochrome c (Cyt-C, non-CMA substrate) in reporter cells treated with A+S for 8, 16, and 24 h. Left panel shows representative immunoblots; right panel shows quantification. Data represent mean ± sd (*n*=3). (**E**) Relative NLuc luminescence in ES2 and SUM159 cells transiently expressing NLuc alone or the HK2-NLuc CMA reporter following CMA activation for up to 24 h. Data represent mean ± sd (*n*=3). *P* values were calculated using two-way ANOVA. (**F**) Immunoblots analysis of lysosomal fractions showing the effect of *LAMP-2A* silencing on lysosomal enrichment of r-HK2, IκBα (CMA substrate), and lysosomal membrane proteins LAMP1 in ES2 cells treated with A+S for 16 h. (**G**) Immunoblot analysis of r-HK2 (detected with anti-HK2 and anti-GFP antibodies) and LAMP-2A expression in ES2 cells following *LAMP-2A* silencing and CMA activation with A+S for 16 h. (**H**) Effect of *LAMP-2A* silencing using two independent siRNAs on relative NLuc luminescence following CMA activation with A+S for 8 and 16 h in ES2 cells. *P* values were calculated using one-way ANOVA. (**I**) Effect of indicated compounds on relative NLuc luminescence in ES2 reporter cells after 24 h treatment. *P* values were calculated using one-way ANOVA. Data points represent individual experiments (*n*=3); bars indicate mean ± sd. Numbers below immunoblot indicate average values from three experiments immunoblots/experiments, normalized to *β*-actin.

In the stable reporter lines, HK2-NLuc (+GFP) was expressed at sub-endogenous levels relative to native HK2, as detected by both HK2 and GFP immunoblotting (Fig. EV1C). Because GFP-tagged reporter exhibited a clear molecular weight shift relative to endogenous HK2, whereas HK2-NLuc co-migrated, we used HK2-GFP-NLuc for immunoblot analyses and HK2-Nluc for luminescence assays. Importantly, the reporter expression did not alter cell growth kinetics, with unchanged doubling times in both cell models (Fig. EV1D), indicating that the reporter is non-perturbative. Average absolute luminescence measurements normalized per 1,000 cells showed no detectable signal in parental cells, even after substrate addition, whereas reporter cells exhibited strong substrate-dependent luminescence (Fig. EV1E), yielding a high signal-to-background ratio suitable for sensitive detection of NLuc signal.

To validate reporter CMA responsiveness, we treated cells with AC220 combined with spautin-1 (A+S), a validated co-treatment known to stimulate CMA in cancer cells (Xia *et al*., 2015). A+S treatment markedly reduced HK2-NLuc protein levels in both stable cell models, as assessed by immunoblotting (Fig. 1B), and this loss was mirrored by a corresponding decrease in NLuc luminescence (Fig. 1C), indicating that the signal reduction reflects reporter degradation. Quantification of luminescence signal using multiple normalization strategies, including average raw relative light units per 1,000 seeded cell, per microgram of total protein, and as percentage of matched vehicle control, yielded comparable reduction with unchanged statistical outcomes (Fig. 1C, EV1F). To reduce well-to-well and plate-to-plate variability, all subsequent luminescence data are therefore presented as percentage of matched vehicle control.

A+S treatment caused a progressive, time-dependent degradation of the HK2-NLuc protein and decline in luminescence signal (Fig. 1D-E). The endogenous CMA substrate mutant p53, decreased with similar kinetics, whereas cytochrome c, a non-CMA substrate, remained unchanged (Fig. 1D). Importantly, a NLuc-reporter not coupled to a CMA substrate was unaffected (Fig. 1E), confirming that signal reduction reflects CMA-mediated degradation rather than a general protein loss or assay drift. Upon CMA activation, both endogenous HK2 and HK2-NLuc accumulated in lysosomal fractions concomitant with increased LAMP-2A abundance (Fig. 1F, EV1G). Depletion of *LAMP-2A* abolished lysosomal enrichment of the reporter and blunted both reporter degradation and luminescence loss (Fig. 1F-H), establishing LAMP-2A dependence of the assay readout.

Finally, we challenged reporter cells with compounds reported to enhance CMA activity, including AR7 (RARα antagonist) (Anguiano *et al*., 2013), geldanamycin (HSP90 inhibitor), and 6-aminonicotinamide (6-AN; G6PD inhibitor) (Finn *et al*, 2005), as well as oxidative stressors, such as paraquat (PQ) and H₂O₂ (Fig. 1I). Most compounds elicited a significant reduction in the NLuc signal consistent with CMA activation. PQ on the other hand, did not reach a statistical significance even though two biological repeats reduced the signal. Although these agents function here as assay benchmarks rather than selective CMA-targeted compounds in cancer, their effects further validate HK2-NLuc as a rapid, reproducible, and quantitative reporter capable of capturing CMA activation across diverse pharmacological and stress-related perturbations.

### Identification of GSK1059615 as a CMA-activating compound

To identify pharmacological activators of CMA that act on upstream, druggable signaling nodes constraining LAMP-2A in cancer, we performed a high-throughput small-molecule screen using the ICCB Known Bioactive Library. This chemically diverse collection is enriched for mechanistically annotated, oncology-relevant compounds, including FDA-approved agents, enabling both pathway inference and rapid progression to *in vivo* proof-of concept studies. Accordingly, each hit could serve both as mechanistic probe of CMA enhancement and as a potential translational lead.

In the primary screen, ES2 reporter cells were treated with 10,523 compounds for 24 h. A+S served as a positive control for CMA activation, DMSO as a negative control, and staurosporine (STS) as a cytotoxicity control. Hits were defined as compounds inducing a ≥50% reduction of NLuc signal relative to vehicle control without overt toxicity (Fig. 2A). Putative hits were then subjected to a secondary screen in both ES2 and SUM159 reporter cells. Compounds that reproducibly reduced NLuc signal in both lines without significantly affecting cell death or proliferation within 24 h were advances (Fig. 2B). This filtering strategy identified GSK1059615 (GSK-615) as a CMA-activating compound. GSK-615 treatment reduced HK2-NLuc reporter abundance in both ES2 and SUM159 cells, as confirmed by immunoblotting using HK2 and GFP antibodies (Fig. 2C). Reporter degradation occurred in a concentration-dependent manner, yielding a sigmoidal response curve with a lower EC₅₀ in SUM159 cells compared to ES2, indicating that CMA activation is achieved at lower GSK-615 concentrations in this cell model (Fig. 2D). Single-dose validation further confirmed significant NLuc signal reduction at 10 µM in ES2 and 5 µM in SUM159 cells (Fig. 2E). Together, these data identify GSK-615 as a dose- and time-dependent compound that promotes degradation of CMA reporter in cancer cells. Importantly, the reduction in luminescence was not attributable to cell loss (Fig. 2F) or decreased reporter mRNA expression (Fig. 2G), indicating that GSK-615 enhance reporter degradation, post-transcriptionally.

**Figure 2.**
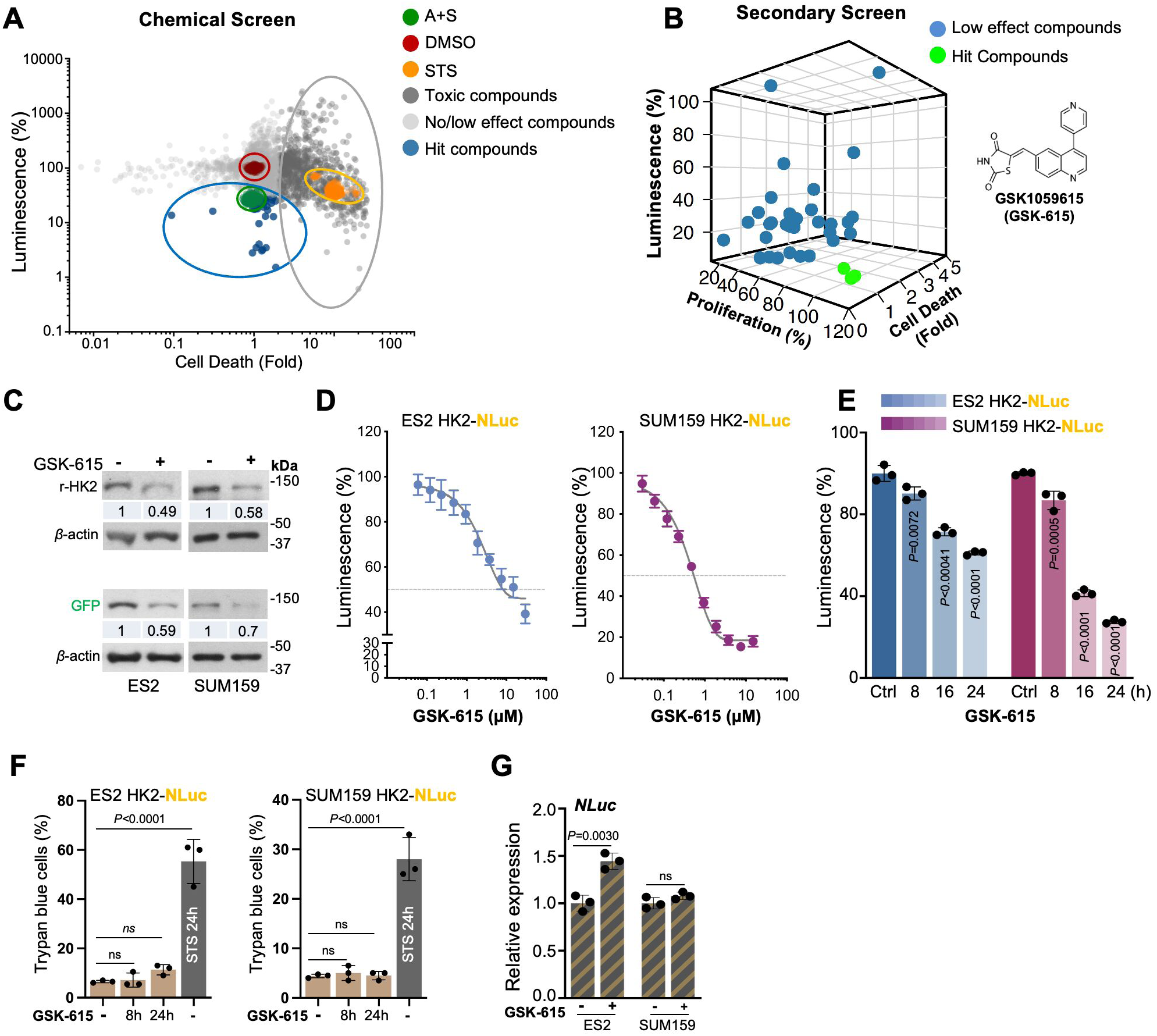
Small-molecule screening identifies GSK-615 as a CMA-activator. (**A**) Scatter plot of the primary chemical screen performed using the ICCB Known Bioactive Collection. A total of 10,523 compounds were screened for effects on HK2-NLuc luminescence (CMA activity) and cell death in ES2 reporter cells. Blue dots indicate hit compounds; red dots indicate negative control (DMSO); green dots indicate positive control for CMA activation (A+S); orange dots indicate the positive control for cytotoxicity (staurosporine; STS). (**B**) Three-dimensional scatter plot showing overlapping hits from secondary screen in ES2 and SUM159 cells performed in duplicate. Top-ranked hits (green) were selected based on reproducible reduction of NLuc signal (CMA activation) without significant effects on proliferation or cell death. (**C**) Immunoblots analysis of reporter-HK2 (r-HK2) degradation in ES2 and SUM159 cells treated with GSK-615 for 24 h, detected using anti-HK2 or anti-GFP antibodies. (**D**) Dose-response curves of GSK-615-induced NLuc decrease in ES2 and SUM159 reporter cells following 24 h treatment. Data represent mean ± sd (*n*=3). Horizontal dashed line indicates EC_50_ values, which represent the concentration required for half-maximal reduction of the CMA reporter signal. (**E**) Relative NLuc luminescence in ES2 and SUM159 reporter cells treated with 10 µM GSK-615 for the indicated times. *P* values were calculated using one-way ANOVA. (**F**) Percentage of trypan blue-positive ES2 and SUM159 cells following GSK-615 treatment up to 24 h. Staurosporine (STS; 24 h) was used as a positive control for cell death. *P* values were calculated using one-way ANOVA. (**G**) Relative NLuc reporter mRNA levels in ES2 and SUM159 cells following GSK-615 treatment for the indicated times. *P* values were calculated using unpaired two-tailed Student’s *t*-test. Data points represent individual experiments (*n*=3); bars indicate mean ± sd. Numbers below immunoblot indicate averages values from three independent experiments relative to control and normalized to *β*-actin.

### GSK-615 activates CMA by upregulating LAMP-2A while sparing macroautophagy

Given that LAMP-2A is the rate limiting determinant of CMA, we examined whether it is affected by GSK-615 and observed a significant increase in LAMP-2A mRNA expression and protein abundance in both ES2 and SUM159 cells (Fig. 3A-B). Consistently, an independent CMA reporter, the photoconvertible PAmCherry-KFERQ (Ho *et al*, 2020), showed an evident increase in lysosomal puncta following GSK-615 treatment, confirming CMA activation at the single-cell level (Fig. 3C). Additionally, GSK-615 reduced canonical CMA substrates, including IκBα and eIF4A1 (Huang & Wang, 2025), and mesenchymal markers, including TGFβR2, Snail or Twist (Zhou *et al*., 2025) (Fig. 3D).

**Figure 3.**
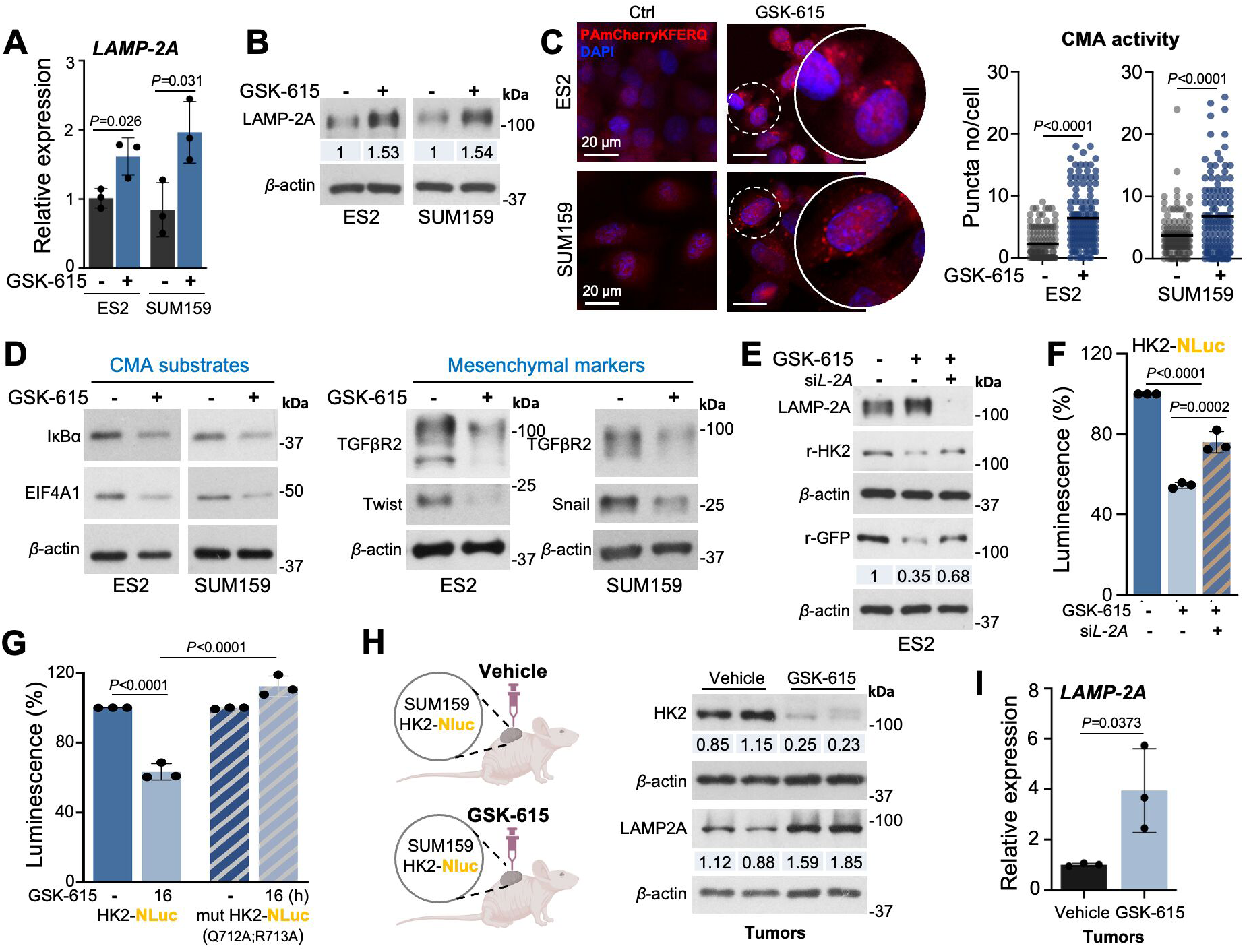
GSK-615 activates CMA through LAMP-2A upregulation without engaging macroautophagy. (**A**) Relative *LAMP-2A* mRNA expression in ES2 and SUM159 cells following GSK1059615 (GSK-615) treatment for 16 h. *P* values were calculated using unpaired two-tailed Student’s *t*-test. (**B**) Immunoblot analysis of LAMP-2A protein levels in ES2 and SUM159 cells treated with GSK-615 for 16 h. (**C**) Confocal microscopy analysis of CMA activity in ES2 and SUM159 cells stably expressing the PAmCherry-KFERQ reporter following GSK-615 treatment for 16 h. Left panels show representative images (DAPI, blue; PAmCherry-KFERQ, red); right panels show quantification of lysosomal puncta per cell. Bars indicate mean. *P* values were calculated using unpaired two-tailed Student’s *t*-test. (**D**) Immunoblot analysis of canonical CMA substrates and mesenchymal markers in ES2 and SUM159 cells treated with GSK-615 for 16 h. (**E**) Immunoblot analysis of LAMP-2A and reporter HK2 (r-HK2; detected with anti-HK2 and anti-GFP antibodies) in ES2 cells treated with GSK-615 for 16 h following *LAMP-2A* silencing. (**F**) Relative NLuc luminescence in ES2 reporter cells treated with GSK-615 for 16 h following *LAMP-2A* silencing. *P* values were calculated using one-way ANOVA. (**G**) Relative NLuc luminescence in ES2 cells expressing wild-type HK2-NLuc or CMA-motif mutant (mut) HK2-NLuc reporters following treatment with 5 µM GSK-615 for 16 h. *P* values were calculated using one-way ANOVA. (**H**) *In vivo* CMA activation by GSK-615. Left, schematic of the experimental design in which SUM159 cells expressing the HK2-GFP-NLuc reporter were xenografted into nude mice and treated with vehicle or GSK-615. Right, representative immunoblots of HK2 and LAMP-2A from tumor lysates. Tumor lysates were analyzed to assess reporter degradation and LAMP-2A induction as readouts of CMA activation *in vivo.* (**I**) Relative *LAMP-2A* mRNA expression in SUM159 xenograft tumors following GSK-615 treatment. Data points represent individual tumors (*n*=3). *P* values were calculated using unpaired two-tailed Student’s *t*-test. Numbers below immunoblot indicate averages from three independent experiments relative to control and normalized to *β*-actin where indicated. Data points represent individual experiments (*n*=3); bars indicate mean ± sd.

Silencing LAMP-2A significantly attenuated GSK-615-induced degradation of HK2-NLuc protein detected using both HK2 and GFP antibodies and partially restored the NLuc signal reduction, whereas *LAMP-2B* or *LAMP1* knockdown had no effect (Fig. 3E-F, EV2A-B). Genetic inhibition of macroautophagy (*ATG5*) or microautophagy (*VPS4A/B*) genes likewise failed to blunt the NLuc signal reduction (Fig. EV2A-B), supporting pathway specificity for GSK-615 on CMA. The incomplete rescue following *LAMP-2A* silencing is likely due to residual lysosomal LAMP-2A and cell-to-cell knockdown heterogeneity. By contrast, a KFERQ-mutant HK2-NLuc reporter, which is incompatible with CMA recognition, did not show luminescence loss upon GSK-615 treatment and exhibited a substantial rescue of NLuc signal compared with the wild type HK2-NLuc reporter (Fig. 3G).

GSK-615 did not induce macroautophagy, as LC3 flux analysis in ES2 or HOS-GFP-LC3 cells revealed no additional LC3-II accumulation with GSK-615 in the presence of lysosomal inhibitors (chloroquine or Leupeptin+NH_4_Cl) beyond the inhibitors alone (Fig. EV2C). In HOS-GFP-LC3 cells, EBSS starvation plus CQ increased LC3 puncta relative to CQ alone, while GSK-615 plus CQ did not (Fig. EV2D). Consistently, transmission electron microscopy revealed no increase in autophagosomes or autolysosomes/amphisomes following GSK-615 treatment, in contrast to AC220, which produced the expected macroautophagy phenotype (Fig. EV2E). Together, these data demonstrate that GSK-615 activates CMA through a LAMP-2A and KFERQ-dependent mechanism across orthogonal readouts, without engaging macroautophagy.

Although GSK-615 is a FDA-approved drug, it is not known whether it can activate CMA an *in vivo* context. To determine this and to assess the utility of the reporter in tumor models, we implanted HK2-GFP-NLuc-expressing SUM159 cells subcutaneously into BALB/c nude mice and treated animals for 14 days with GSK-615 (25 mg/kg, i.p.) or vehicle (Fig. 3H). Tumor lysates from GSK-615 treated mice showed increased *LAMP-2A* mRNA and protein abundance together with reduced HK2-NLuc reporter protein (Fig. 3H-I). These findings mirror the cell-based assays and indicate that GSK-615 engages CMA in tumors, with the reporter serving as a tumor-context readout of CMA activity, establishing GSK-615 as a tool compound to probe CMA regulation *in vitro* and *in vivo*.

### Genome-wide CRISPR screen identifies MEK-ERK as a brake on CMA

In parallel with the compound screen, we performed a genome-wide CRISPR-Cas9 loss-of-function screen to identify genetic regulators of CMA in cancer cells. Cas9-expressing ES2 cells stably carrying the HK2-GFP-NLuc reporter were transduced with a pooled sgRNA library targeting ∼20,000 genes (∼80,000 sgRNAs). After 10 days to allow gene knockout and protein turnover, cells were sorted based on reporter GFP intensity into GFP-high (CMA unaffected or inhibited) and GFP-low (reporter degraded/ CMA activated) populations (Fig. 4A). Because loss of a negative regulator of CMA results in enhanced reporter degradation, sgRNAs enriched in the GFP-low fraction identify genes that normally function as brakes on CMA. Genomic DNA from each fraction was sequenced and analyzed at the sgRNA level by Internal Replicate Analysis (IRA). Genes were scored based on (i) statistical significance (*P* < 0.05) for enrichment in GFP-low versus GFP-high populations, (ii) ≥ 1.5-fold enrichment, and (iii) representation by at least three independent sgRNA exclusively in GFP-low fraction (Fig. 4B). HK2 itself ranked among the top hits (Fig. 4B), consistent with direct loss of the reporter backbone, serving as an internal positive control for screening performance.

**Figure 4.**
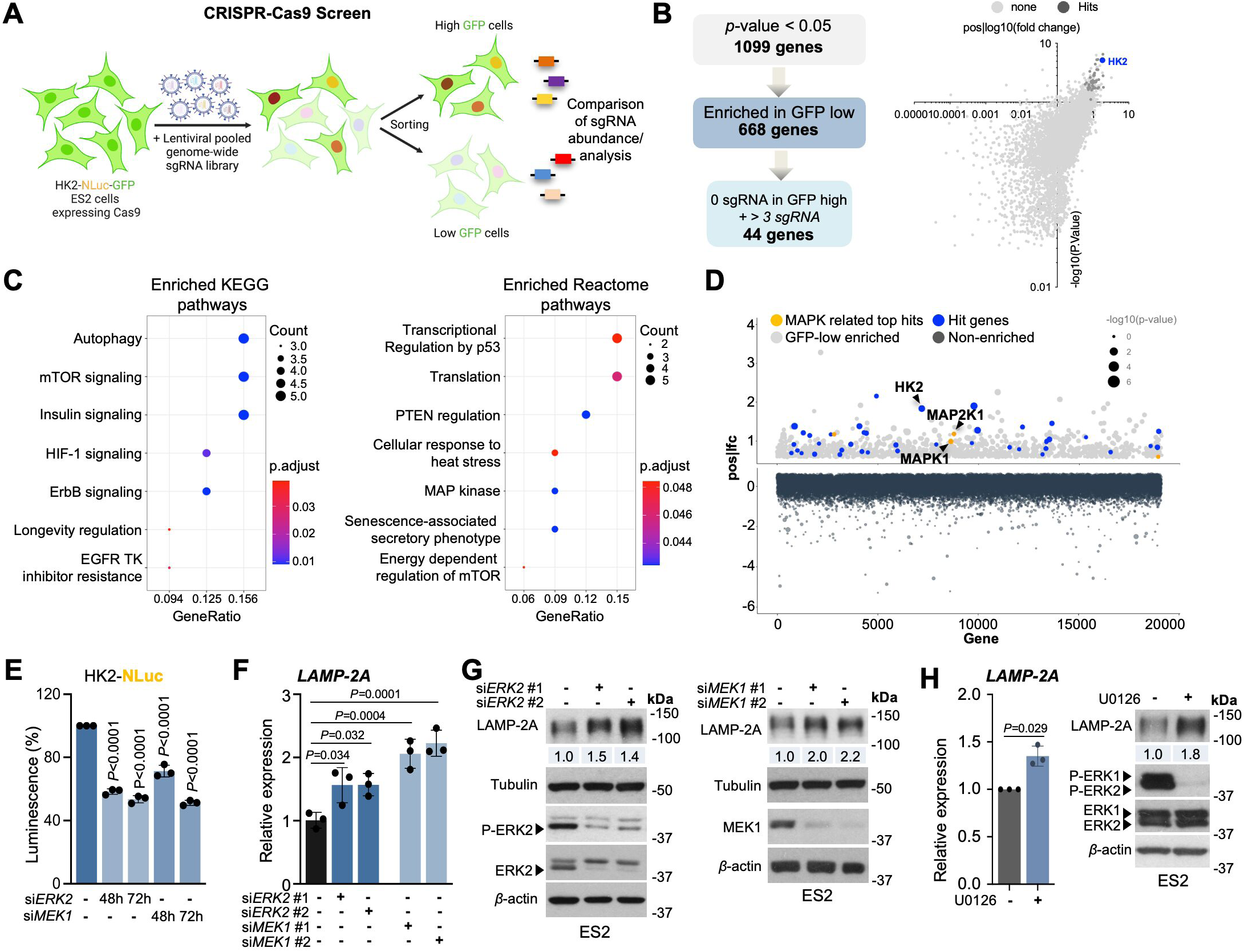
Genome-wide CRISPR screening identifies MEK-ERK signaling as a negative regulator of CMA. (**A**) Schematic of the genome-wide CRISPR–Cas9 loss-of-function screen using the HK2-GFP-NLuc CMA reporter in ES2 cells. Cells were transduced with a pooled sgRNA library, selected for 10 days, sorted into GFP-low (CMA-activated) and GFP-high (CMA-unaffected) populations, and sgRNA representation was analyzed by next-generation sequencing. (**B**) Hit selection strategy. Left, Venn diagram illustrating cutoff criteria applied to identify candidate CMA repressors. Right, volcano plot showing gene-level enrichment based on log₂ fold change (x-axis) and statistical significance (-log₁₀ *P* value; y-axis). Dark gray dots indicate hit genes; HK2 is highlighted in blue as an internal control. (**C**) KEGG and Reactome pathway enrichment analysis of hit genes identified in the CRISPR screen. (**D**) Scatter plot showing –log₁₀ *P* values for all genes identified in the screen. Dark gray dots represent genes not enriched in the GFP-low population (log₂ fold change ≤ 0); light gray dots represent genes enriched in GFP-low cells but not meeting full hit criteria; blue dots indicate hit genes; orange dots highlight MAPK pathway–associated hits. **(E)** Relative NLuc luminescence and (**F**) relative *LAMP-2A* mRNA expression in ES2 cells following 48 h knockdown of ERK2 or MEK1 using two independent siRNAs. *P* values were calculated using one-way ANOVA. (**G**) Immunoblot analysis of LAMP-2A and total and phosphorylated ERK1/2 following 48 h ERK2 depletion (left), and LAMP-2A and total and phosphorylated MEK1 following 48 h MEK1 depletion (right). (**H**) Relative *LAMP-2A* mRNA levels (left) and immunoblot analysis of LAMP-2A, total ERK1/2, and phosphorylated ERK1/2 (right) in ES2 cells following 48 h treatment with the MEK inhibitor U0126. *P* values were calculated using unpaired two-tailed Student’s *t*-test. Numbers below immunoblot indicate averages from three independent experiments, normalized to *β*-actin.

Pathway enrichment analysis (KEGG and Reactome) on the hit set recovered known CMA-linked processes, including insulin/PI3K-AKT, mTOR, and HIF1/oxidative stress signaling (Arias *et al*., 2015; Endicott *et al*., 2020; Hubbi *et al*, 2013). Remarkably, the analysis highlighted ErbB-MAP kinase cascades as a previously uncharacterized negative regulator of CMA (Fig. 4C), nominating MAP2K1 (MEK1) and its substrate MAPK1 (ERK2) as candidate key players (Fig. 4D). We validated this axis genetically using *MEK1* or *ERK2* siRNA and observed that individual knockdown of these genes alone was able to reduce the HK2-NLuc signal, consistent with increased CMA activity (Fig. 4E), along with increased *LAMP-2A* mRNA and protein levels in ES2 cells (Fig. 4F-G). We next tested a selective, non-ATP-competitive MEK1/2 inhibitor, U0126, that suppresses ERK phosphorylation and downstream signaling outputs. In ES2 cells U0126 treatment increased *LAMP-2A* mRNA and protein levels (Fig. 4H), mirroring the genetic data, supporting a MEK-ERK-imposed transcriptional brake on *LAMP-2A* expression.

### Oncogenic wiring stratifies LAMP-2A and predicts CMA activation routes

We next asked whether baseline MAPK-related oncogenic alterations in epithelial (A549) and mesenchymal (SUM159 and ES2) cancer cells correlate with LAMP-2A abundance. These cell lines harbor distinct driver landscapes that differentially tune MEK-ERK and PI3K-AKT signaling (Fig. 5A). ES2 carries a BRAF alteration that constitutively activates the MEK-ERK pathway, whereas A549 is a KRAS mutant and SUM159 harbors HRAS-associated alterations with engagement of the PI3K-axis. Immunoblotting of cell lysates showed that SUM159 cells displayed the highest P-AKT (S473/T308), with P-ERK levels comparable to A549 but lower than ES2, which displayed the highest P-ERK and the lowest P-AKT levels (Fig. 5B). Consistent with these differences in phosphorylation patterns, LAMP-2A protein abundance was lowest in ES2 and highest in A549, with SUM159 showing intermediate levels, mirroring baseline differences in CMA activity across these lines (Fig. 5C). By contrast, *LAMP-2A* transcript levels were similar between SUM159 and A549 (Fig. 5D). Given that KRAS/HRAS activates ERK less potently than BRAF (Brandt *et al*, 2019; Hymowitz & Malek, 2018), the transcript-protein discordance in SUM159 is consistent with post-transcriptional suppression of LAMP-2A via AKT-GFAP-dependent control of stability, whereas the ERK-dominant ES2 phenotype supports MEK-ERK mediated repression of *LAMP-2A* transcription. Together these patterns predict that MEK/ERK inhibition should more effectively activate CMA in ES2 cells, whereas PI3K/AKT inhibition should preferentially increase CMA in SUM159.

**Figure 5.**
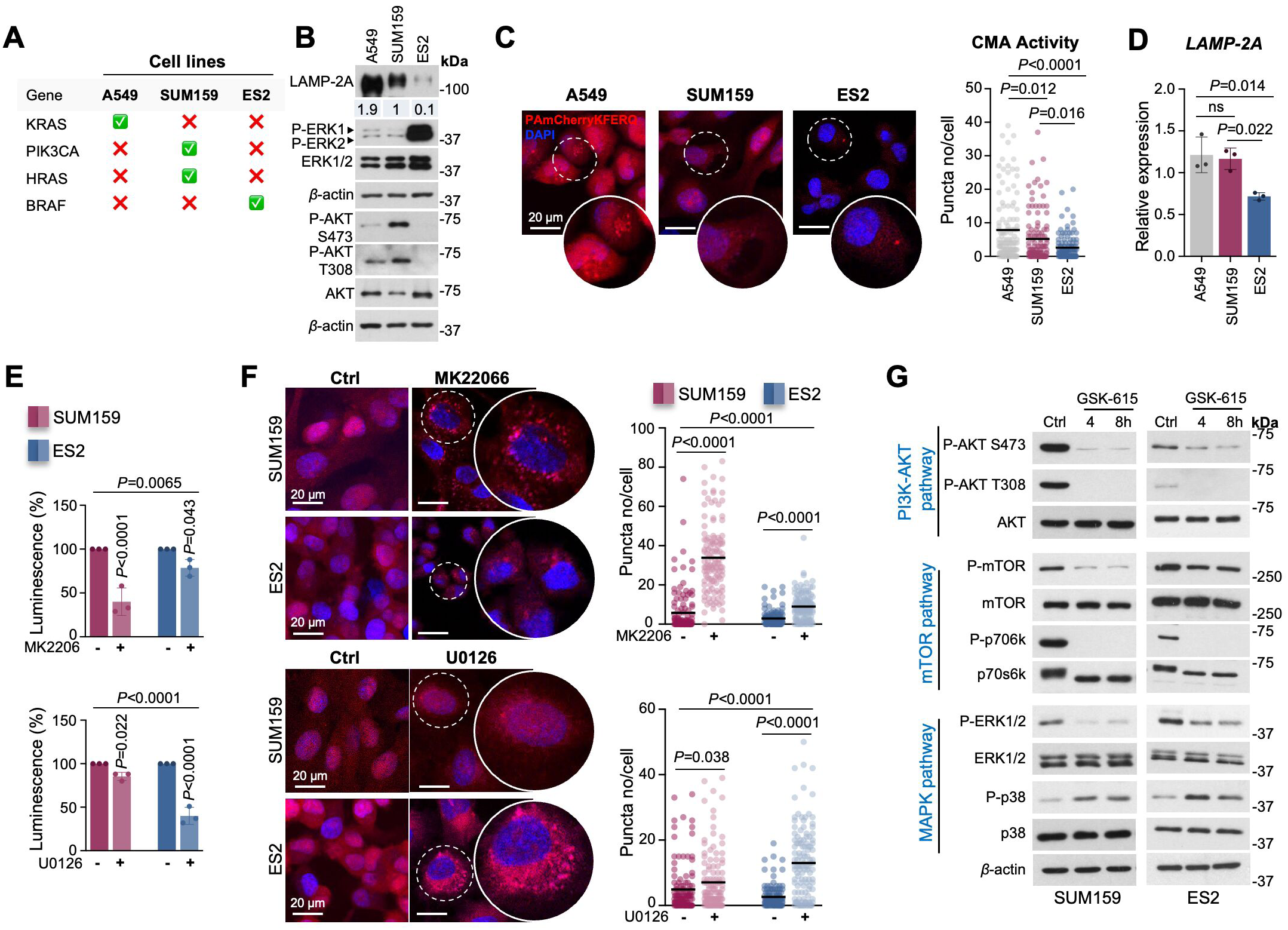
Oncogenic signaling context determined routes to CMA activation. (**A**) Table summarizing oncogenic driver alterations in A549, SUM159, and ES2 cells. **(B)** Immunoblot analysis of LAMP-2A, total and phosphorylated ERK1/2, and AKT in epithelial (A549) and mesenchymal (ES2, SUM159) cancer cells. Numbers below blots indicate averages from three independent experiments, normalized to *β*-actin and expressed relative to A549. **(C)** Confocal microscopy of A549, ES2, and SUM159 cells stably expressing the PAmCherry-KFERQ CMA reporter. Left, representative images showing DAPI (blue) and PAmCherry-KFERQ (red). Right, quantification of CMA puncta per cell. Bars represent mean; *P* values were calculated using one-way ANOVA. (**D**) Relative *LAMP-2A* mRNA expression in A549, ES2, and SUM159 cells. Data points represent individual experiments (*n*=3); bars indicate mean ± sd. *P* values were calculated using one-way ANOVA. (**E**) Relative NLuc activity in ES2 and SUM159 cells stably expressing the HK2-NLuc CMA reporter following 48 h treatment with the AKT inhibitor MK2206 (top) or the MEK inhibitor U0126 (bottom). Data points represent individual experiments (*n*=3); bars indicate mean ± sd. *P* values were calculated using two-way ANOVA. (**F**) Confocal microscopy of ES2 and SUM159 cells stably expressing PAmCherry-KFERQ following 48 h treatment with MK2206 (top) or U0126 (bottom). Left, representative images; right, quantification of puncta per cell. Bars represent mean; *P* values were calculated using two-way ANOVA. (**G**) Immunoblot analysis of total and phosphorylated ERK1/2, p38, AKT, mTOR, and p70S6K following GSK1059615 treatment.

To test these predictions, we quantified CMA activity using both HK2-NLuc reporter and PAmCherry-KFERQ assays, comparing U0126 and the pan-AKT inhibitor (MK2206) on ES2 and SUM159 cells. U0126 treatment resulted in a greater HK2-NLuc signal decrease, indicative of higher CMA activity, and a more pronounced PAmCherry-KFERQ puncta compared with the pan-AKT inhibitor in ES2 cells. In contrast, MK2206 elicited stronger CMA activation in SUM159, exceeding the effect of U0126 (Fig. 5E-F, EV3A). These results indicate that ERK inhibition preferentially restores CMA in ERK-dependent ES2 cells, whereas relief of AKT signaling more effectively activates CMA in SUM159, consistent with their baseline oncogenic background.

Finally, we examined the signaling effects of GSK-615 in both ES2 and SUM159 cells, where GSK-615 treatment reduced levels of P-AKT (S473, T308), P-mTOR, and downstream signaling (P-S6K) without altering the total protein levels, confirming effective suppression of the PI3K-AKT-mTOR axis (Fig. 5G). In parallel, GSK-615 decreased P-ERK1/2 (T202/Y204) while increasing p38 MAPK phosphorylation (T180/Y182), indicating coordinated attenuation of ERK signaling together with p38 engagement. This signaling profile corresponds to conditions that favor both relief of *LAMP-2A* transcription and stabilization of LAMP-2A at lysosomes, consistent with the observed activation of CMA in reporter assays following GSK-615 treatment. Notably, because the EC₅₀ reflects CMA reporter responsiveness rather than general drug sensitivity, the lower EC₅₀ observed in SUM159 (Fig. 2D), indicates that CMA activation is achieved at lower GSK-615 concentrations in this model. This aligns with PI3K-AKT signaling functioning as the dominant suppressive input on CMA in SUM159 cells and with GSK-615 acting as a potent PI3K pathway inhibitor, such that AKT inhibition alone is sufficient to relieve the lysosomal LAMP-2A stability brake. By contrast, ES2 cells rely more heavily on ERK-dependent transcriptional repression of *LAMP-2A*, which may require greater ERK attenuation to surpass the threshold for transcriptional de-repression.

### Transcription Factors of the Forkhead Family contribute to *LAMP-2A* expression

To identify transcriptional drivers linking oncogenic pathway relief to *LAMP-2A* induction, we performed RNA-seq in ES2 and SUM159 cells treated with GSK-615 compared to control, reasoning that CMA activation by GSK-615 involves transcriptional factor (TF)-mediated *LAMP-2A* expression. TF activity was inferred with ISMARA (Integrated System for Motif Activity Response Analysis), which integrates motif occurrence with gene expression changes to estimate motif activity scores. ISMARA analysis revealed extensive transcriptional reprogramming upon GSK-615 treatment, with multiple transcriptional programs showing fluctuating activity in both cell lines (Fig 6A). Comparative analysis across ES2 and SUM159 cells identified a shared set of TFs whose inferred activities were consistently altered by GSK-615 (Fig. 6B). Ranking these factors by cumulative activity change highlighted Forkhead box (FOX) family members, among the most strongly increased programs (Fig. 6C). FOX factors are known targets of ERK-mediated inhibitory phosphorylation, suggesting a mechanistic link between ERK attenuation and FOX factors activation (Boccitto & Kalb, 2011; Farhan *et al*, 2017). In contrast, transcriptional programs centered in MYC/MXI1 and CREB1 that are ERK-proximal and reliant on shared CBP/p300 co-activators (Sears *et al*, 2000; Wang *et al*, 2018), were among the most strongly decreased, pointing to a coordinated transcriptional shift favoring FOXO-dependent gene expression. Consistent with this model, heatmap visualization of ISMARA-predicted target genes revealed concurrent upregulation of FOXO1/FOXP1-associated transcriptional programs and downregulation of CREB/MYC-driven targets following GSK-615 treatment (Fig. 6D). Inferred FOXO1 and FOXP1 motif activity correlated positively with their transcript abundance (TPM), consistent with increased expression contributing to elevated transcriptional output following GSK-615 treatment (Fig. 6E). qPCR and immunoblot analyses confirmed induction of FOXO1 and FOXP1 at both mRNA and protein levels in ES2 and SUM159 cells following GSK-615 treatment, with cell line-specific differences in magnitude but concordant directionality across the assays (Fig. 6F-G).

**Figure 6.**
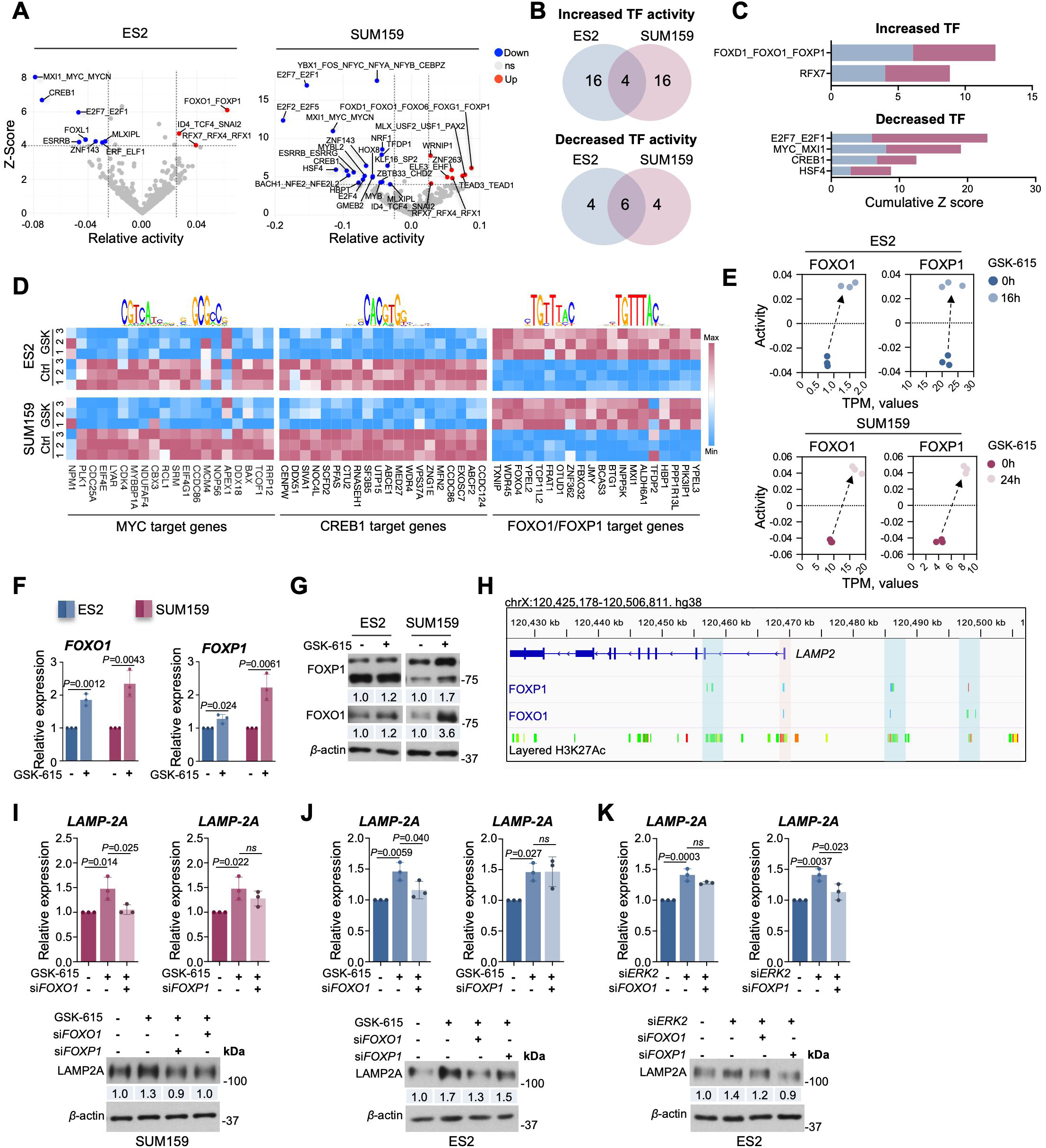
FOXO1 and FOXP1 promote *LAMP-2A* transcription downstream of ERK inhibition. (**A**) Volcano plots of ISMARA-inferred transcription factor (TF) activity in ES2 and SUM159 cells treated with GSK-615. TFs with significantly increased activity are highlighted in blue, decreased activity in red, and non-significant changes (P > 0.05 and <2-fold change) in grey. (**B**) Venn diagram showing shared and cell line-specific TFs among the top upregulated and downregulated factors following GSK-615 treatment. (**C**) Cumulative ISMARA Z-scores of shared TFs upregulated or downregulated in both ES2 and SUM159 cells, highlighting FOX-family enrichment among induced programs. (**D**) Heat maps of RNA-seq expression changes for predicted FOXO1/FOXP1, MYC, and CREB1 target gene sets in ES2 and SUM159 cells treated with GSK-615 for 16 h. Sequence logos shown above each heat map represent the corresponding transcription factor DNA-binding motifs used by ISMARA to define target gene sets; two distinct logos are shown for FOXO1 (left) and FOXP1 (right). (**E**) Relationship between ISMARA-inferred motif activity (y-axis) and transcript abundance (TPM; x-axis) for FOXO1 and FOXP1 in ES2 and SUM159 cells following GSK-615 treatment. (**F**) Relative *FOXO1* and *FOXP1* mRNA expression in ES2 and SUM159 cells after 16 h of GSK-615 treatment. Data points represent individual experiments (*n*=3); bars show mean ± sd. *P* values were calculated using unpaired two-tailed Student’s *t*-test. (**G**) Immunoblot analysis of FOXO1 and FOXP1 protein levels in ES2 and SUM159 cells treated with GSK-615 for 16 h. (**H**) IGV genome browser view showing FOXP1, FOXO1, and H3K27ac ChIP-seq enrichment across the human *LAMP2* locus (hg38), compiled from ChIP-Atlas datasets across multiple cell types (collapsed view). Shaded regions indicate putative promoter (red) and enhancer (blue) elements. (**I**) Relative *LAMP-2A* mRNA (upper panels) and protein levels (lower panels) in SUM159 cells treated with GSK-615 for 16 h alone or combined with *FOXO1* or *FOXP1* siRNA-mediated knockdown for 72 h. (**J**) Relative LAMP-2A mRNA (upper panels) and protein levels (lower panel) in ES2 cells treated with GSK-615 for 16 h or combined with FOXO1 or FOXP1 siRNA-mediated knockdown for 72 h. (**K**) Relative *LAMP-2A* mRNA (upper panels) and protein levels (lower panels) in ES2 cells following *ERK2* knockdown alone or combined with *FOXO1* or *FOXP1* siRNA-mediated knockdown for 48 h. Control and GSK-615 conditions are shown for independent ANOVA comparisons. Data points represent individual experiments (*n*=3); bars show mean ± sd. *P* values were calculated using unpaired one-way ANOVA. Numbers below immunoblot indicate averages from three independent experiments, normalized to *β*-actin where indicated.

We next examined publicly available ChIP-seq datasets for FOXO1 and FOXP1 occupancy across the human *LAMP2* locus. This analysis revealed enrichment of FOXO1 and FOXP1 at promoter-proximal and multiple distal regulatory enhancer regions of *LAMP2* that overlap with H3K27ac-marked active chromatin (Fig. 6H), supporting a direct transcriptional input of these transcription factors into *LAMP2* gene regulation.

To functionally test the contribution of FOXO1 and FOXP1 to LAMP-2A induction, we performed siRNA-mediated knockdown experiments. Silencing *FOXO1* significantly attenuated the GSK-615-induced increase in *LAMP-2A* mRNA and protein levels in both ES2 and SUM159 cells (Fig. 6I-J, EV3B). *FOXP1* knockdown produced a similar directional effect but did not reach statistical significance under these conditions. In ES2 cells, ERK2 depletion uncovered a selective dependence on FOXP1 for maximal *LAMP-2A* induction, whereas FOXO1 silencing attenuated *LAMP-2A* expression independently of ERK status (Fig. 6K, EV3C). Together, these perturbations demonstrate selective modulation of *LAMP-2A* expression by FOXO1 and FOXP1 in cancer cells.

Collectively these data identify FOXO1 and FOXP1 as mediators linking ERK pathway attenuation to transcriptional induction of *LAMP-2A*. This transcriptional program provides a mechanistic bridge between oncogenic signaling relief and restoration of CMA capacity, positioning FOXO1 activity as a determinant of *LAMP-2A* expression and CMA competence in cancer cells.

## Discussion

This study identifies MEK–ERK signaling as a dominant, reversible brake on chaperone-mediated autophagy (CMA) in cancer and establishes LAMP-2A as a key point of convergence for oncogenic signaling control of this pathway. By combining chemical and genome-scale genetic screening, we show that sustained MAPK activity suppresses CMA at the level of its rate-limiting receptor and that pharmacological relief of this constraint restores CMA activity in cancer cells and xenografts without engaging macroautophagy.

Our findings place ERK signaling upstream of CMA regulation through direct control of *LAMP-2A* expression. In support of this model, a recent study reported that the clinically approved MEK1 inhibitor trametinib promotes CMA-dependent turnover of lipogenic enzymes in mouse liver, demonstrating that MEK inhibition is permissive for CMA activity *in vivo* (Chen *et al*, 2025). That study, however, was performed in a non-malignant context and did not address transcriptional regulation of CMA components. Our work extends these observations by showing that, in human cancer cells, ERK signaling actively represses *LAMP-2A* transcription and that relief of this repression represents a principal mechanism by which CMA can be restored. Together, these findings establish MEK-ERK as a conserved but context-dependent regulator of CMA, acting through distinct mechanistic layers in physiological versus oncogenic settings. At the transcriptional level, we identify Forkhead family transcription factors as key mediators linking ERK and AKT relief to LAMP-2A induction. Transcriptomic and motif activity analyses reveal a coordinated shift toward FOX-driven gene programs accompanied by suppression of ERK-proximal CREB and MYC activity (Sears *et al*., 2000; Wang *et al*., 2018). Functional perturbation supports a primary role for FOXO1, with a contributory role for FOXP1, in driving *LAMP-2A* transcription. These results are consistent with established negative regulation of FOX factors by ERK and AKT signaling (Farhan *et al*., 2017; Laissue, 2019; Roy *et al*, 2010; Xie *et al*, 2012) and suggest that CMA repression in ERK-high cancers reflects constrained FOXO activity rather than an intrinsic incapacity to activate the pathway. Our data further support an indirect role for CREB and MYC, likely mediated through competition for shared transcriptional co-activators (Wang *et al*, 2020) rather than direct repression of the *LAMP-2* locus.

More broadly, our results indicate that CMA output in cancer cells is not a fixed property but an integrated consequence of convergent oncogenic signaling. ERK signaling functions as a dominant transcriptional brake on *LAMP-2A* in contexts of sustained MAPK activity, while AKT–mTORC2 signaling imposes an additional post-translational constraint on LAMP-2A stability (Arias *et al*., 2015). In contrast, p38 activity acts permissively by stabilizing LAMP-2A at the lysosomal membrane (Li *et al*., 2017). GSK-615 engages all three axes simultaneously, providing a mechanistic explanation for the magnitude and robustness of CMA activation observed across distinct tumor contexts. This integrative model also clarifies why prior interventions targeting single pathways, such as PI3K/AKT inhibition or stress induction, can elevate CMA without defining a clear transcriptional control point (Endicott *et al*., 2020; Li *et al*., 2017).

Importantly, these findings link ERK-mediated CMA repression to the TGFβ-driven mesenchymal program characteristic of advanced tumors (Derynck *et al*, 2021). ERK signaling is a central driver of tumor progression and therapy resistance (Guo *et al*, 2020; Sulzmaier & Ramos, 2013; Timofeev *et al*, 2024), and our model predicts that ERK-high cancers exist in a CMA-low state due to suppressed *LAMP-2A* expression. This provides a mechanistic connection to our previous observation that CMA loss promotes dedifferentiation and metastatic competence in mesenchymal tumors (Zhou *et al*., 2025). We propose that ERK-dependent suppression of CMA sustains TGFβ signaling by limiting turnover of EMT-associated effectors, while TGFβ signaling further reinforces ERK activity, forming a feed-forward loop that stabilizes the invasive mesenchymal state. Restoration of CMA disrupts this loop, reducing TGFβ output and attenuating mesenchymal traits.

Conceptually, this work reframes CMA from a passive stress-responsive pathway to a regulated and druggable output of oncogenic signaling. In tumors characterized by high ERK activity and mesenchymal features, restoring CMA represents a rational strategy to dampen pro-invasive signaling networks. This framework yields testable predictions: ERK-high tumors should exhibit lower basal CMA, increased sensitivity to CMA restoration upon MAPK pathway inhibition, and CMA-dependent clearance of EMT-related effectors. Together, our findings establish ERK-imposed repression of LAMP-2A as a reversible control point for CMA and highlight pathway-aligned CMA restoration as a context-specific strategy to counteract aggressive tumor states.

## Methods

### Cell culture and Treatments

ES2 and A549 cells were cultured in RPMI medium supplemented with 10% (v/v) heat-inactivated fetal bovine serum (FBS), 100 U/mL penicillin, 100 U/mL streptomycin, and 1% (w/v) glutamine. SUM159 cells were maintained in Ham’s F12 medium supplemented with 5% (v/v) heat-inactivated FBS, 100 U/mL penicillin, 100 U/mL streptomycin, 5 µg/mL insulin, and 1 µg/mL hydrocortisone. HOS GFP-LC3 cells were cultured in DMEM supplemented with 10% (v/v) heat-inactivated FBS, 100 U/mL penicillin, 100 U/mL streptomycin, and 1% (w/v) glutamine. All cell lines were maintained at 37°C in a humidified incubator with 5% CO₂ and used in logarithmic growth phase. Cells were treated with spautin-1 (10 μM), AC220 (1.5 μM), geldanamycin (4 μM), 6-aminonicotinamide (6-AN; 10 μM), paraquat (PQ; 2.5 mM), AR7 (40 μM), H₂O₂ (250 μM), U0126 (10 μM), MK2206 (1 μM), or GSK1059615 (5 μM for SUM159 and 10 μM for ES2 cells, unless otherwise specified). Staurosporine (STS; 5 μM) was used as a positive control for cytotoxicity. Lysosomal inhibition was performed using leupeptin (40 μM), NH₄Cl (10 mM), or chloroquine (50 μM). Macroautophagy was induced by incubation in Earle’s Balanced Salt Solution (EBSS).

### Generation of HK2-Nluc CMA reporter cell lines

The coding sequence of wild-type (WT) human hexokinase 2 (HK2) and a CMA motif–deficient mutant (Q712A/R713A), with or without an N-terminal green fluorescent protein (GFP) tag, were cloned into the pNLF1-HIF1A vector (see Reagent and Tools table) using NEBuilder HiFi DNA Assembly (New England Biolabs). WT and mutant HK2 coding sequences were PCR-amplified using primers HK2Fw and HK2Rv, while GFP-tagged constructs were amplified using primers HK2Fw and HK2GFPRv (Appendix Table S1) from the FLHKII-pGFPN3 plasmid (Sun *et al*, 2008). The pNLF1-HIF1A backbone was digested with NheI and XhoI to remove the HIF1A insert prior to assembly. The resulting pNLF1-HK2 constructs contained the NLF1 promoter, HK2 (WT or mutant), GFP (where applicable), NanoLuc (NLuc), a stop codon, and a polyadenylation signal. All constructs were sequence-verified prior to use. Stable cell lines expressing either HK2-GFP-NLuc or HK2-NLuc were generated by transfection using ViaFect reagent (Promega), according to the manufacturer’s instructions. Forty-eight hours after transfection, cells were selected with G418 (800 μg/mL) for 7 days. Single-cell clones were isolated by limiting dilution in 96-well plates and maintained under selection for an additional two weeks. Individual clones were screened for luminescence activity using the Nano-Glo Luciferase assay. Throughout the study, HK2-GFP-NLuc expressing cells were used for immunoblotting to allow discrimination of the reporter fusion protein from endogenous HK2, whereas HK2-NLuc expressing cells were used for luminescence-based assays to avoid potential interference from GFP.

### Nano-Glo Luciferase assay

Cells stably expressing the CMA reporter were seeded in 96-well plates at subconfluent density 24 h prior to treatment. Following treatment, Nano-Glo® Luciferase assays (Promega) were performed according to the manufacturer’s instructions. Briefly, culture medium was replaced with 20 μL of fresh medium per well, followed by addition of 20 μL of reconstituted Nano-Glo reagent (substrate diluted 1:50 in lysis buffer). Luminescence was measured using a GloMax® microplate reader (Promega).

### Cell death assay

Cells were treated with GSK-615 and harvested at 8 and 24 h post-treatment. Cell death was assessed by trypan blue exclusion using 0.4% trypan blue solution. Trypan blue-positive cells were quantified and expressed as a percentage of total cells. Staurosporine (STS; 5 μM) was used as a positive control.

### Cell proliferation assay

Cells were seeded in triplicate in 96-well plates. Cell viability was assessed at 24, 48, and 72 h using the CellTiter-Glo Luminescent Cell Viability Assay (Promega), according to the manufacturer’s instructions. Equal volumes of CellTiter-Glo reagent were added to each well, and luminescence was measured using a GloMax® microplate reader (Promega). Viability values were normalized to the corresponding 0 h measurement for each cell line.

### Compound screening

ES2 reporter cells (1,500 cells per well) were seeded in 384-well plates in a final volume of 40 μL using an automated dispenser. After 24 h, compounds from the ICCB Known Bioactive Libraries (10,542 compounds, including FDA-approved drugs; ICCB-Longwood Screening Facility, Harvard Medical School, Boston, MA, USA) were transferred by pin-transfer robot (100 nL per well), yielding final compound concentrations of 10-30 μM in a total volume of 35 μL per well. DMSO concentration (0.3%) was equalized across all wells, including controls. CMA activity was assessed 24 h after compound addition using the Nano-Glo Luciferase assay. To monitor cytotoxicity, 15 μL of culture medium from each well was transferred to a separate 384-well plate prior to lysis, and cell death was quantified using the ToxiLight BioAssay. No edge effects were detected. The primary screen was performed in duplicate. Screening data were analyzed using Z scores, calculated as Z=(X−Ave_neg)/SD_neg. Compounds inducing ≥50% reduction in luminescence relative to DMSO controls and <1.2-fold increase in cell death were considered primary hits. Hits were subsequently validated in a secondary screen performed in duplicate using both ES2 and SUM159 reporter cells. Secondary screening criteria included a 40-60% reduction in luminescence, <1.2-fold (SUM159) or <2-fold (ES2) increase in cell death, and preserved cell proliferation. For proliferation measurements, nuclei were stained with Hoechst 33342, imaged using an ImageXpress Micro Widefield Confocal System, and quantified by automated image analysis. DMSO served as a negative control, spautin-1 (10 μM) combined with AC220 (1.5 μM) as a positive control for CMA activation, and staurosporine (STS; 5 μM) as a positive control for cytotoxicity.

### CRISPR-Cas9 screening

ES2 cells stably expressing the HK2-GFP CMA reporter and Cas9 nuclease were generated as previously described (Schmierer *et al*, 2017). Briefly, cells were transduced with a lentiviral construct encoding Cas9, blasticidin resistance, and a single guide RNA (gRNA) targeting *HPRT1*. Functional Cas9 activity was verified by sequential selection with blasticidin and 6-thioguanine, a nucleotide analog lethal to *HPRT1*-proficient cells. The genome-wide Brunello sgRNA library (Doench *et al*, 2016) was cloned and packaged into lentivirus as previously described. Viral titer was determined by serial dilution in ES2 HK2-GFP cells followed by puromycin selection. Cas9-expressing ES2 HK2-GFP cells were transduced with the Brunello library at a multiplicity of infection (MOI) of approximately 0.4, corresponding to ∼1,000 cells per sgRNA, in the presence of polybrene (2 μg/mL). Transduced cells were selected with puromycin (1 μg/mL) from day 2 to day 4 post-transduction and subsequently split into two independent biological replicates. Cells were cultured until day 10 post-transduction, maintaining a minimum coverage of 40 × 10⁶ cells per replicate throughout the experiment. At day 10, cells were sorted based on reporter GFP intensity into GFP-high (CMA inactive or repressed) and GFP-low (CMA activated) populations using a BD FACSJazz cell sorter. For each condition, 7 × 10⁶ cells were collected per replicate. Genomic DNA was isolated using the QIAamp DNA Blood Maxi Kit (Qiagen), and sgRNA sequences were PCR-amplified as previously described. Sequencing was performed at the National Genomics Infrastructure (SciLifeLab) using an Illumina HiSeq 2500 platform. Next-generation sequencing data were analyzed using the MAGeCK pipeline (Li *et al*, 2014). Strictly standardized mean difference (SSMD) scores were calculated for each sgRNA and averaged to obtain gene-level scores. Candidate genes were selected based on statistical significance (P < 0.05), enrichment in the GFP-low population, and representation by at least three independent sgRNAs. Pathway enrichment analysis was performed on the top 44 ranked genes using the clusterProfiler package (v 4.6.2) (Wu *et al*, 2021) with hypergeometric testing for KEGG and Reactome pathways. Multiple testing correction was applied using the Benjamini–Hochberg method, with adjusted P < 0.05 considered significant.

### siRNA transfection

Cells were transfected with siRNA (20–60 nM final concentration) using INTERFERin (Polyplus) according to the manufacturer’s instructions. Knockdown efficiency was assessed 48–72 h post-transfection by quantitative PCR and immunoblotting. Sequences of all siRNAs used in this study are provided in Reagent and Tools table.

### RNA extraction and qPCR

Total RNA was extracted using the PureLink RNA Mini Kit (Thermo Fisher Scientific) with on-column DNase digestion according to the manufacturer’s instructions. Complementary DNA (cDNA) was synthesized using the iScript cDNA Synthesis Kit (Bio-Rad). Quantitative PCR (qPCR) was performed using Maxima SYBR Green Master Mix (Thermo Fisher Scientific) on a StepOnePlus Real-Time PCR System (Applied Biosystems), using approximately 20 ng cDNA per reaction. Primer sequences are listed in Appendix Table S1. Relative gene expression was calculated using the ΔΔCT method, with ACTB, GAPDH, TUBA1A, ESD, or HPRT serving as reference genes, as indicated.

### Western blotting

Cells were lysed in RIPA buffer (150 mM NaCl, 1% NP-40, 1% sodium deoxycholate, 0.1% SDS, 25 mM Tris–HCl pH 7.6) supplemented with protease and phosphatase inhibitor cocktails. Protein concentration was determined using the BCA assay. Equal amounts of protein (10–30 μg) were resolved by SDS–PAGE and transferred onto nitrocellulose membranes. Membranes were immunoblotted with the indicated primary and secondary antibodies (Appendix Table S2). Immunoreactive bands were detected using enhanced chemiluminescence (ECL) and visualized on X-ray film (Fujifilm).

### Animal studies

All animal experiments were approved by the Stockholm Animal Ethics Committee (ethical permit no. N116/16) and conducted in accordance with Swedish animal welfare regulations. Five-week-old female BALB/c nude mice (Balb/cAnN-Foxn1^nu/nu^, Janvier Labs) were housed under pathogen-free conditions at constant temperature (21°C) and humidity (50-60%), with ad libitum access to food and water. SUM159 cells stably expressing HK2-GFP-NLuc (5 × 10⁶ cells in 100 μL of 50% Matrigel in PBS) were injected subcutaneously into the flank, with two injection sites per mouse (n=2 mice per group). Beginning 3 days post-injection, mice were treated with either vehicle (5% glucose solution) or GSK1059615 (25 mg/kg) by intraperitoneal injection every second day. Mice were euthanized at day 14 and tumors were immediately processed for RNA and protein extraction.

### Lysosomal fractionation

Lysosomal fractionation was performed using the Lysosome Isolation Kit (Sigma-Aldrich) as previously described (Hao *et al*., 2019; Kacal & Vakifahmetoglu-Norberg, 2022). Briefly, cells were homogenized in 1× extraction buffer by 15 strokes with a 25-gauge syringe needle. Post-nuclear supernatants were cleared by differential centrifugation, normalized for protein content, and layered onto an OptiPrep density gradient (8-27%; Sigma-Aldrich). Samples were subjected to ultracentrifugation, and gradient fractions were collected, pelleted, and resuspended. Fraction purity was assessed by immunoblotting for lysosomal markers (LAMP1, LAMP-2A) and the absence or depletion of markers for other organelles, including CTSD, TOM40, LDHA, HSC/= and LC3B, as shown in Expanded View Figure 1 in (Zhou *et al*., 2025). Fractions enriched for lysosomal markers were used for subsequent analyses.

### High throughput quantification of GFP-LC3 puncta

HOS GFP-LC3 cells were seeded in 96-well plates and treated as indicated. Cells were washed with PBS and incubated in L-15 medium supplemented with 10% (v/v) fetal bovine serum and 1% (w/v) glutamine. Nuclei were stained with Hoechst 33342 to enable cell segmentation. Images were acquired using an ImageXpress Micro Widefield Confocal System (Molecular Devices). Automated image analysis was performed using CellProfiler software. Nuclei were identified based on Hoechst 33342 signal, and cytoplasmic regions were defined by subtracting nuclear masks from the GFP-LC3 channel. GFP-LC3 puncta were detected using intensity-based thresholding, and the number of puncta per cell was quantified for each condition in an automated, unbiased manner.

### PAmCherry-KFERQ CMA reporter assay

Lentiviral particles were generated by co-transfecting HEK293FT cells with the pSIN-PAmCherry-KFERQ-NE reporter plasmid (Ho *et al*., 2020) together with ViraPower Lentiviral Expression System packaging plasmids (pLP1, pLP2, and pLP/VSVG), according to the manufacturer’s instructions. Target cells were transduced with viral supernatant in the presence of polybrene and selected with puromycin 48 h post-transduction for two weeks. For CMA analysis, reporter-expressing cells were seeded on coverslips and treated as indicated. Photoactivation of PAmCherry-KFERQ was performed by exposure to 360–390 nm UV light to convert the reporter to its fluorescent form. Cells were subsequently fixed in 4% (w/v) paraformaldehyde for 15 min at room temperature and mounted using Vectashield mounting medium containing DAPI (Vector Laboratories). Images were acquired using a Zeiss LSM900 confocal microscope. CMA activity was quantified by measuring the number of lysosomal PAmCherry-KFERQ puncta per cell using FIJI (ImageJ) in an automated and blinded manner (n=40 cells per replicate per condition).

### Transmission electron microscopy (TEM)

Transmission electron microscopy was performed at the Electron Microscopy Unit (EMIL), Karolinska Institutet. ES2 cells were fixed in 2.5% (v/v) glutaraldehyde in 0.1 M phosphate buffer (pH 7.4), rinsed in the same buffer, and post-fixed with 2% (w/v) osmium tetroxide in 0.1 M phosphate buffer for 2 h at 4°C. Samples were dehydrated through a graded ethanol series followed by acetone and embedded in LX-112 epoxy resin. Ultrathin sections (∼80 nm) were cut using a Leica EM UC7 ultramicrotome, collected on copper grids, and contrasted with uranyl acetate followed by Reynolds lead citrate. Sections were examined using a Hitachi HT7700 transmission electron microscope, and digital images were acquired with a Veleta CCD camera. Autophagic structures, including autophagosomes and autolysosomes, were quantified manually in a blinded manner from randomly selected fields.

### RNA-Sequencing and Transcription Factor Activity prediction by ISMARA

Total RNA was isolated from control- and GSK1059615-treated cells and submitted for bulk RNA sequencing at the National Genomics Infrastructure (NGI), SciLifeLab, Karolinska Institutet. Library preparation and sequencing were performed according to standard NGI protocols. Differential gene expression analysis was conducted using DESeq2 in R (version 4.0.2). Genes with an adjusted *P* value < 0.05 were considered significantly differentially expressed unless otherwise stated. Transcription factor (TF) activity profiles were inferred using the Integrated System for Motif Activity Response Analysis (ISMARA) algorithm, which integrates promoter motif occurrence with expression changes to estimate motif activity (*z*-scores). ISMARA analysis was performed using FASTQ files as input, as described previously (Balwierz *et al*, 2014).

### ChIP-seq data analysis

FOXP1, FOXO1, and H3K27ac ChIP-seq data were retrieved from ChipAtlas (hg38 assembly). Datasets included GSM3633494 (FOXP1), GSM5214708, GSM4971622, and GSM2136846 (FOXO1). Significance thresholds were set at 100 for FOXP1/FOXO1 peaks across all cell types and 500 for H3K27ac peaks. Genome browser tracks at the LAMP2 locus were generated using the Integrative Genomics Viewer (IGV) in collapsed format for FOXP1/FOXO1 and layered format for H3K27ac.

### Statistics

Data are presented as mean ± standard deviation (sd) from at least three independent biological experiments unless otherwise indicated. Statistical analyses were performed using GraphPad Prism (version 9). Comparisons between two groups were assessed using unpaired two-tailed Student’s *t*-tests. Comparisons involving more than two groups were analyzed using one-way analysis of variance (ANOVA) followed by Tukey’s multiple-comparison test. Differences in treatment effects between cell lines were analyzed using two-way ANOVA. Statistical significance was defined as *P* < 0.05. Exact sample sizes, statistical tests, and *P* values are reported in the figure legends.

## Acknowledgements

We thank Dr. Eeva-Liisa Eskelinen (University of Turku, Finland) for evaluation of transmission electron microscopy images; Dr. Paulo Jannig (Karolinska Institutet, Sweden) for assistance with ChIP analysis; Dr. Gerald McInerney (Karolinska Institutet, Sweden) for providing the HOS GFP-LC3 cells. We thank Dr. Yuqing Hao for assistance with lysosomal fractionation. We acknowledge Dr. Amanda T. Ouchida for generating stable cell lines and contributing to early stages of the study, including initial validation of the CMA reporter, execution of the chemical screen, and preparation of samples for the CRISPR-Cas9 screen. We also thank Yashar Esmaeilian and Daniel R. Vaz for experimental assistance, and Ece Kilinc, Davide Chiesi, and Aironas Los (project and master’s students) for technical support. This work was supported by the Ragnar Söderberg Foundation, and the Swedish Cancer Society (25 4481 Pj).

## Author contribution

Eva Berenger: Data curation; Formal analysis; Validation; Investigation; Visualization; Writing and editing. Merve Kacal: Data curation; Formal analysis; Validation; Investigation; Methodology. Elena Kochetkova: Data curation; Formal analysis; Validation; Investigation. Alice Maestri: Data curation; Formal analysis; Investigation; Visualization. Boxi Zhang: Software; Formal analysis; Visualization. Suresh Sajwan: Methodology. Mattias Mannervik: Resources; Methodology. Vitaliy O. Kaminskyy: Conceptualization; Supervision; Data curation; Formal analysis; Validation; Investigation; Methodology; Writing and review. Erik Norberg: Resources; Supervision; Funding acquisition; Validation; Investigation. Helin Vakifahmetoglu-Norberg: Conceptualization; Resources; Formal analysis; Supervision; Funding acquisition; Methodology; Project administration; Writing and editing.

## Disclosure and competing interest statement

The authors declare no conflict of interest.

## Data Availability

Proteomics data were deposited in ProteomeXchange via the PRIDE database platform with accession number PXD058467. UTR for the MS data:

https://www.ebi.ac.uk/pride/archive/projects/PXD058467

RNA sequencing data were deposited in the ArrayExpress platform with accession number E-MTAB-14665. UTR for the RNAseq data:

https://www.ebi.ac.uk/biostudies/arrayexpress/studies/E-MTAB-14665?query=E-MTAB-14665

## Expanded View Figure legends

**Figure EV1. Validation of NLuc-based reporters for quantitative CMA detection.** (**A**) Schematic illustration of NLuc fusion reporter constructs (HK2-GFP-NLuc and HK2-NLuc) used to generate stable CMA reporter cell lines. (**B**) Immunoblot analysis of LAMP-2A expression in normal lung fibroblasts (WI-38, MRC-5) and cancer cell lines (A549, SUM159, ES2). Numbers below immunoblot indicate averages from three independent experiments, normalized to *β*-actin where indicated. (**C**) Immunoblot detection of HK2-GFP-NLuc reporter expression in ES2 and SUM159 cells using anti-HK2 (r-HK2) or anti-GFP antibodies. Numbers below immunoblot indicate averages from three independent experiments: upper values correspond to the reporter and lower values to endogenous HK2, normalized to endogenous HK2. (**D**) Cell proliferation analysis of ES2 and SUM159 cells with or without HK2-GFP-NLuc reporter expression. Data represent mean ± sd from *n*=3 independent experiments. *P* values were calculated using two-way ANOVA. (**E**) Luminescence measurements normalized to 1,000 cells in parental ES2 and SUM159 cells and in cells expressing the HK2-NLuc reporter, measured in the presence or absence of luciferase substrate. (**F**) Raw luminescence signal (left) and luminescence normalized to total protein content (right) in ES2 and SUM159 cells expressing HK2-NLuc following 16 h treatment with A+S. (**G**) Immunoblot analysis of reporter HK2 (r-HK2), endogenous HK2, IκBα (CMA substrate), LAMP-2A, and LAMP1 in whole-cell lysates (input) and isolated lysosomal fractions following 16 h treatment with A+S. LAMP1 serves as a lysosomal marker. Data points represent individual experiments (*n*=3); bars indicate mean ± sd. *P* values were calculated using unpaired two-tailed Student’s *t*-test.

**Figure EV2. GSK-615 activates CMA without inducing macroautophagy.** (**A**) Immunoblot analysis of LAMP-2A, LAMP-2B, LAMP1, ATG5, and VPS4A/B in ES2 cells following 48 h siRNA-mediated knockdown of the indicated genes. (**B**) Relative NLuc luminescence in ES2 cells expressing the HK2-NLuc reporter 48 h after knockdown of *LAMP-2A* or *LAMP1* (left) and *VPS34* or *ATG5* (right), followed by GSK-615 treatment for 16 h. Data points represent individual experiments (*n*=3); bars indicate mean ± sd. (**C**) Immunoblot analysis of LC3-I and LC3-II in ES2 cells (upper panel) and HOS GFP-LC3 reporter cells (lower panel) treated with GSK-615 in the presence or absence of lysosomal inhibitors (chloroquine or leupeptin + NH₄Cl) for 16 h. *β*-actin serves as a loading control. (**D**) Representative confocal images of GFP-LC3–expressing HOS cells treated as indicated for 16 h (left) and quantification of GFP-LC3 puncta per cell (right). Box plots show median and 5-95 percentiles: >100 cells per condition. (**E**) Representative transmission electron microscopy images (left) and quantification of autolysosomes per cell (right) in ES2 cells treated for 16 h with AC220 or GSK-615. Box plots indicate median and 10-90 percentiles: *n*=40 cells per condition. *m*, mitochondria; *N*, nucleus; *AL*, autolysosome; *L*, lysosome. *P* values were calculated using unpaired one-way ANOVA.

**Figure EV3**. **Modulation of ERK and AKT signaling and Forkhead transcription factors.** (**A**) Immunoblot analysis of total and phosphorylated ERK and AKT in ES2 and SUM159 cells treated for 48 h with the MEK inhibitor U0126 or the AKT inhibitor MK2206. (**B**) Immunoblot analysis of FOXP1 and FOXO1 protein levels in SUM159 (left) and ES2 (right) cells following 16 h GSK-615 treatment, with or without 48 h *FOXP1* or *FOXO1* siRNA-mediated knockdown. (**C**) Immunoblot analysis of ERK2, FOXP1, and FOXO1 protein levels in ES2 cells following 48 h ERK2 knockdown alone or in combination with *FOXP1* or *FOXO1* knockdown for 48 h.

## Notes

### Competing Interest Statement

The authors have declared no competing interest.

## References

Anguiano J, Garner TP, Mahalingam M, Das BC, Gavathiotis E, Cuervo AM (2013) Chemical modulation of chaperone-mediated autophagy by retinoic acid derivatives. In: Nat Chem Biol, pp. 374–382. United States

Arias E, Cuervo AM (2020) Pros and Cons of Chaperone-Mediated Autophagy in Cancer Biology. Trends Endocrinol Metab 31: 53–66

Arias E, Koga H, Diaz A, Mocholi E, Patel B, Cuervo AM (2015) Lysosomal mTORC2/PHLPP1/Akt Regulate Chaperone-Mediated Autophagy. Mol Cell 59: 270–284

Balwierz PJ, Pachkov M, Arnold P, Gruber AJ, Zavolan M, van Nimwegen E (2014) ISMARA: automated modeling of genomic signals as a democracy of regulatory motifs. In: Genome Res, pp. 869–884. United States

Boccitto M, Kalb RG (2011) Regulation of Foxo-dependent transcription by post-translational modifications. In: Curr Drug Targets, pp. 1303–1310. United Arab Emirates

Bourdenx M, Martin-Segura A, Scrivo A, Rodriguez-Navarro JA, Kaushik S, Tasset I, Diaz A, Storm NJ, Xin Q, Juste YR et al (2021) Chaperone-mediated autophagy prevents collapse of the neuronal metastable proteome. Cell 184: 2696–2714 e2625

Brandt R, Sell T, Luthen M, Uhlitz F, Klinger B, Riemer P, Giesecke-Thiel C, Schulze S, El-Shimy IA, Kunkel D et al (2019) Cell type-dependent differential activation of ERK by oncogenic KRAS in colon cancer and intestinal epithelium. In: Nat Commun, p. 2919. England

Chen J, Berg J, Burns CM, Jia H, Li X, Miller RA, Endicott SJ, Garcia G (2025) The lifespan-extending MEK1 inhibitor trametinib promotes regulation of de novo lipogenesis enzymes by chaperone-mediated autophagy. In: Front Aging, p. 1621808. Switzerland

Chen R, Li P, Fu Y, Wu Z, Xu L, Wang J, Chen S, Yang M, Peng B, Yang Y et al (2023) Chaperone-mediated autophagy promotes breast cancer angiogenesis via regulation of aerobic glycolysis. In: PLoS One, p. e0281577. United States

Derynck R, Turley SJ, Akhurst RJ (2021) TGFbeta biology in cancer progression and immunotherapy. In: Nat Rev Clin Oncol, pp. 9–34. England

Ding ZB, Fu XT, Shi YH, Zhou J, Peng YF, Liu WR, Shi GM, Gao Q, Wang XY, Song K et al (2016) Lamp2a is required for tumor growth and promotes tumor recurrence of hepatocellular carcinoma. Int J Oncol 49: 2367–2376

Doench JG, Fusi N, Sullender M, Hegde M, Vaimberg EW, Donovan KF, Smith I, Tothova Z, Wilen C, Orchard R et al (2016) Optimized sgRNA design to maximize activity and minimize off-target effects of CRISPR-Cas9. Nat Biotechnol 34: 184–191

Endicott SJ, Ziemba ZJ, Beckmann LJ, Boynton DN, Miller RA (2020) Inhibition of class I PI3K enhances chaperone-mediated autophagy. In: J Cell Biol, United States

Farhan M, Wang H, Gaur U, Little PJ, Xu J, Zheng W (2017) FOXO Signaling Pathways as Therapeutic Targets in Cancer. In: Int J Biol Sci, pp. 815–827. Australia

Finn PF, Mesires NT, Vine M, Dice JF (2005) Effects of small molecules on chaperone-mediated autophagy. In: Autophagy, pp. 141–145. United States

Gomes LR, Menck CFM, Cuervo AM (2017) Chaperone-mediated autophagy prevents cellular transformation by regulating MYC proteasomal degradation. In: Autophagy, pp. 928–940. United States

Gomez-Sintes R, Xin Q, Jimenez-Loygorri JI, McCabe M, Diaz A, Garner TP, Cotto-Rios XM, Wu Y, Dong S, Reynolds CA et al (2022) Targeting retinoic acid receptor alpha-corepressor interaction activates chaperone-mediated autophagy and protects against retinal degeneration. In: Nat Commun, p. 4220. England

Guo YJ, Pan WW, Liu SB, Shen ZF, Xu Y, Hu LL (2020) ERK/MAPK signalling pathway and tumorigenesis. In: Exp Ther Med, pp. 1997–2007. Greece

Hao Y, Kacal M, Ouchida AT, Zhang B, Norberg E, Vakifahmetoglu-Norberg H (2019) Targetome analysis of chaperone-mediated autophagy in cancer cells. In: Autophagy, pp. 1558–1571. United States

Ho PW, Leung CT, Liu H, Pang SY, Lam CS, Xian J, Li L, Kung MH, Ramsden DB, Ho SL (2020) Age-dependent accumulation of oligomeric SNCA/alpha-synuclein from impaired degradation in mutant LRRK2 knockin mouse model of Parkinson disease: role for therapeutic activation of chaperone-mediated autophagy (CMA). In: Autophagy, pp. 347–370. United States

Huang J, Wang J (2025) Selective protein degradation through chaperone-mediated autophagy: Implications for cellular homeostasis and disease (Review). Mol Med Rep 31

Huang YB, Tian LL, Zhu ZW, Zhou KG, Lai X, Peng YZ, Wu Z, Tong WF, Wang H, Wang XJ et al (2025) Apigenin enhances Nrf2-induced chaperone-mediated autophagy and mitigates alpha-synuclein pathology: Implications for Parkinson’s disease therapy. Phytomedicine 141: 156652

Hubbi ME, Hu H, Kshitiz, Ahmed I, Levchenko A, Semenza GL (2013) Chaperone-mediated autophagy targets hypoxia-inducible factor-1alpha (HIF-1alpha) for lysosomal degradation. In: J Biol Chem, pp. 10703–10714. United States

Hubert V, Weiss S, Rees AJ, Kain R (2022) Modulating Chaperone-Mediated Autophagy and Its Clinical Applications in Cancer. In: Cells, Switzerland

Hymowitz SG, Malek S (2018) Targeting the MAPK Pathway in RAS Mutant Cancers. In: Cold Spring Harb Perspect Med, United States

Ikami Y, Terasawa K, Watabe T, Yokoyama S, Hara-Yokoyama M (2022) The two-domain architecture of LAMP2A within the lysosomal lumen regulates its interaction with HSPA8/Hsc70. In: Autophagy Rep, pp. 205–209. United States

Kacal M, Vakifahmetoglu-Norberg H (2022) Isolation of Autophagy Competent Lysosomes from Cancer Cells by Differential Large-Scale Multilayered Density Gradient Centrifugations. Methods Mol Biol 2445: 27–38

Kacal M, Zhang B, Hao Y, Norberg E, Vakifahmetoglu-Norberg H (2021) Quantitative proteomic analysis of temporal lysosomal proteome and the impact of the KFERQ-like motif and LAMP2A in lysosomal targeting. In: Autophagy, pp. 3865–3874. United States

Kaushik S, Cuervo AM (2018) The coming of age of chaperone-mediated autophagy. In: Nat Rev Mol Cell Biol, pp. 365–381. England

Kirchner P, Bourdenx M, Madrigal-Matute J, Tiano S, Diaz A, Bartholdy BA, Will B, Cuervo AM (2019) Proteome-wide analysis of chaperone-mediated autophagy targeting motifs. In: PLoS Biol, p. e3000301. United States

Laissue P (2019) The forkhead-box family of transcription factors: key molecular players in colorectal cancer pathogenesis. In: Mol Cancer, p. 5. England

Li W, Xu H, Xiao T, Cong L, Love MI, Zhang F, Irizarry RA, Liu JS, Brown M, Liu XS (2014) MAGeCK enables robust identification of essential genes from genome-scale CRISPR/Cas9 knockout screens. In: Genome Biol, p. 554. England

Li W, Zhu J, Dou J, She H, Tao K, Xu H, Yang Q, Mao Z (2017) Phosphorylation of LAMP2A by p38 MAPK couples ER stress to chaperone-mediated autophagy. In: Nat Commun, p. 1763. England

Li Y, Sheng M, Li W, Liu S, Wang B, Liu B, Luo M, Zhou X, Xia Q, Hong S et al (2025) Targeting chaperone-mediated autophagy inhibits properties of glioblastoma stem cells and restores anti-tumor immunity. In: Nat Commun, p. 440. England

Liu J, Wang L, He H, Liu Y, Jiang Y, Yang J (2023) The Complex Role of Chaperone-Mediated Autophagy in Cancer Diseases. In: Biomedicines, Switzerland

Motomura K, Ueda E, Boateng A, Sugiura M, Kadoyama K, Hitora-Imamura N, Kurauchi Y, Katsuki H, Seki T (2024) Identification of a novel aromatic-turmerone analog that activates chaperone-mediated autophagy through the persistent activation of p38. In: Front Cell Dev Biol, p. 1418296. Switzerland

Pajares M, Rojo AI, Arias E, Diaz-Carretero A, Cuervo AM, Cuadrado A (2018) Transcription factor NFE2L2/NRF2 modulates chaperone-mediated autophagy through the regulation of LAMP2A. In: Autophagy, pp. 1310–1322. United States

Qiao L, Hu J, Qiu X, Wang C, Peng J, Zhang C, Zhang M, Lu H, Chen W (2023) LAMP2A, LAMP2B and LAMP2C: similar structures, divergent roles. In: Autophagy, pp. 2837–2852. United States

Rios J, Sequeida A, Albornoz A, Budini M (2020) Chaperone Mediated Autophagy Substrates and Components in Cancer. Front Oncol 10: 614677

Roy SK, Srivastava RK, Shankar S (2010) Inhibition of PI3K/AKT and MAPK/ERK pathways causes activation of FOXO transcription factor, leading to cell cycle arrest and apoptosis in pancreatic cancer. In: J Mol Signal, p. 10. England

Saberiyan M, Gholami S, Ejlalidiz M, Rezaeian Manshadi M, Noorabadi P, Hamblin MR (2025) The dual role of chaperone-mediated autophagy in the response and resistance to cancer immunotherapy. Crit Rev Oncol Hematol 210: 104700

Saha T (2012) LAMP2A overexpression in breast tumors promotes cancer cell survival via chaperone-mediated autophagy. In: Autophagy, pp. 1643–1656. United States

Schmierer B, Botla SK, Zhang J, Turunen M, Kivioja T, Taipale J (2017) CRISPR/Cas9 screening using unique molecular identifiers. In: Mol Syst Biol, p. 945. Germany

Sears R, Nuckolls F, Haura E, Taya Y, Tamai K, Nevins JR (2000) Multiple Ras-dependent phosphorylation pathways regulate Myc protein stability. Genes Dev 14: 2501–2514

Seike T, Terasawa K, Iwata T, Guan JL, Watabe T, Yokoyama S, Hara-Yokoyama M (2024) Site-specific photo-crosslinking of Hsc70 with the KFERQ pentapeptide motif in a chaperone-mediated autophagy and microautophagy substrate in mammalian cells. Biochem Biophys Res Commun 736: 150515

Sulzmaier FJ, Ramos JW (2013) RSK isoforms in cancer cell invasion and metastasis. In: Cancer Res, pp. 6099–6105. United States

Sun L, Shukair S, Naik TJ, Moazed F, Ardehali H (2008) Glucose phosphorylation and mitochondrial binding are required for the protective effects of hexokinases I and II. In: Mol Cell Biol, pp. 1007–1017. United States

Timofeev O, Giron P, Lawo S, Pichler M, Noeparast M (2024) ERK pathway agonism for cancer therapy: evidence, insights, and a target discovery framework. In: NPJ Precis Oncol, p. 70. England

Ueda E, Ohta T, Konno A, Hirai H, Kurauchi Y, Katsuki H, Seki T (2022) D-Cysteine Activates Chaperone-Mediated Autophagy in Cerebellar Purkinje Cells via the Generation of Hydrogen Sulfide and Nrf2 Activation. In: Cells, Switzerland

Vakifahmetoglu-Norberg H, Kim M, Xia HG, Iwanicki MP, Ofengeim D, Coloff JL, Pan L, Ince TA, Kroemer G, Brugge JS et al (2013) Chaperone-mediated autophagy degrades mutant p53. In: Genes Dev, pp. 1718–1730. United States

Valdor R, Mocholi E, Botbol Y, Guerrero-Ros I, Chandra D, Koga H, Gravekamp C, Cuervo AM, Macian F (2014) Chaperone-mediated autophagy regulates T cell responses through targeted degradation of negative regulators of T cell activation. In: Nat Immunol, pp. 1046–1054. United States

Wang D, Wang Y, Zou X, Shi Y, Liu Q, Huyan T, Su J, Wang Q, Zhang F, Li X et al (2020) FOXO1 inhibition prevents renal ischemia-reperfusion injury via cAMP-response element binding protein/PPAR-gamma coactivator-1alpha-mediated mitochondrial biogenesis. In: Br J Pharmacol, pp. 432–448. England

Wang H, Xu J, Lazarovici P, Quirion R, Zheng W (2018) cAMP Response Element-Binding Protein (CREB): A Possible Signaling Molecule Link in the Pathophysiology of Schizophrenia. Front Mol Neurosci 11: 255

Wu T, Hu E, Xu S, Chen M, Guo P, Dai Z, Feng T, Zhou L, Tang W, Zhan L et al (2021) clusterProfiler 4.0: A universal enrichment tool for interpreting omics data. In: Innovation (Camb), p. 100141. United States

Xia HG, Najafov A, Geng J, Galan-Acosta L, Han X, Guo Y, Shan B, Zhang Y, Norberg E, Zhang T et al (2015) Degradation of HK2 by chaperone-mediated autophagy promotes metabolic catastrophe and cell death. In: J Cell Biol, pp. 705–716. United States

Xie L, Ushmorov A, Leithauser F, Guan H, Steidl C, Farbinger J, Pelzer C, Vogel MJ, Maier HJ, Gascoyne RD et al (2012) FOXO1 is a tumor suppressor in classical Hodgkin lymphoma. In: Blood, pp. 3503–3511. United States

Yan J, Liu D, Wang J, You W, Yang W, Yan S, He W (2024) Rewiring chaperone-mediated autophagy in cancer by a prion-like chemical inducer of proximity to counteract adaptive immune resistance. Drug Resist Updat 73: 101037

Yao R, Shen J (2023) Chaperone-mediated autophagy: Molecular mechanisms, biological functions, and diseases. In: MedComm (2020), p. e347. China

Zhou X, Berenger E, Shi Y, Shirokova V, Kochetkova E, Becirovic T, Zhang B, Kaminskyy VO, Esmaeilian Y, Hosaka K et al (2025) Chaperone-mediated autophagy regulates the metastatic state of mesenchymal tumors. In: EMBO Mol Med, pp. 747–774. Germany

